# Effects of a Sequential Application of Plant Protection Products on Soil Microbes and Free-Living Nematodes in a Field Experiment

**DOI:** 10.1101/2025.02.17.638612

**Authors:** Camilla Drocco, Saúl Fernandes, Liyan Xie, Marion Devers, Bernhard Förster, Fabrice Martin-Laurent, Sana Romdhane, Aymé Spor, Clémence Thiour-Mauprivez, Anja Coors

## Abstract

During crop growth cycle, often several different plant protection products (PPPs) are applied, in combination or sequentially. Such sequential applications result in unintentional mixtures of residues that may affect ecosystem services supported by non-target organisms such as soil microbes and nematodes. This scenario of sequential PPP application is frequent in agricultural practice but rarely addressed experimentally at field scale with regard to environmental impacts. The objective of this study was to evaluate the effect of individual and sequential application of three PPPs (the herbicide clopyralid, the insecticide zeta-cypermethrin, and the fungicide pyraclostrobin) on soil microbial communities, and on the abundance of free-living nematode. Single applications (at 1× or 10× the agronomical dose) were made to triplicated field plots with each one of the PPPs or all three PPPs in sequence, with untreated plots serving as controls. Plots were sampled prior to each application, and 7 and 28 days thereafter. The fungal community’s composition and abundance were found to be more susceptible than the bacterial community to PPP applications, while the bacterial was mainly driven by the in-field heterogeneity of soil properties. Transient effects of PPP applications were detected on nematode abundance. Higher tier ecotoxicological studies offer greater ecological relevance compared to the standard laboratory tests but are challenged by environmental variations that should be accounted for when evaluating the ecotoxicity of pesticides on soil organisms.

## INTRODUCTION

Soil is a complex substrate that hosts a great biological diversity, which is vital to keep soil healthy and resilient. The interplay between diverse communities is crucial as they provide mutual benefits to each other (Wall et al. 2012) and support soil health. The intensive use of agricultural lands, including mechanical perturbations and agrochemicals application, raises concern about the long-term soil communities functioning and resilience (Murray et al. 2006; Bardgett and van der Putten 2014; Bender et al. 2016). Indeed, agrochemicals residues are frequently detected in agricultural soils (FAO and UNEP 2021; Vieira et al. 2023) and can impair ecological functions provided by microbes (Storck et al. 2018; Gallego et al. 2019; Romdhane et al. 2019) such as nutrient cycles, carbon sequestration and disease suppression (Ward and Jensen, 2014; Singh 2015; Nannipieri et al. 2017; Schloter et al. 2018). The environmental risk assessment (ERA) of plant protection products (PPPs) and their active ingredients (a.i) require ecotoxicity tests on various standard model organisms. For soil microbiota, however, only functional endpoints measured in the nitrogen transformation test (OECD 216) are required, which may not cover possible impacts on all soil microbiota’s ecological functions (Karpouzas et al. 2022).

Nematodes, the most abundant metazoans in soil (Boufahja et al. 2011), occur worldwide and interact with a broad range of organisms including microorganisms (van den Hoogen et al. 2019; Neilson et al. 2020; van den Hoogen et al. 2020). They have been extensively utilized for soil health monitoring purposes. Nematodes are not represented among the standard test organisms considered in an ERA, despite evidence that they may be susceptible to adverse PPPs effects (Zhao et al. 2013; Waldo et al. 2019; Grabau et al. 2020; Höss et al. 2022). The European Food and Safety Authority (EFSA) proposed a range of possible endpoints, including microorganisms and nematodes, that could cover identified shortcomings in the ERA of PPPs for in-soil organisms (EFSA Panel on Plant Protection Products and their Residues (PPR) et al. 2017).

The ERA assesses prospectively the individual risk of PPPs and their a.i. after their intentional release into the environment and the risk of mixtures (combined exposures) as detailed in a guidance document (EFSA, 2019). However, PPPs are frequently applied sequentially during the crop growth cycle, which may impact the ability of the soil communities to recover from previous treatments. Effects of such unintended (coincidental) mixtures of different PPP residues resulting from sequential (serial) applications on the soil compartment remain largely unknown (Weisner et al. 2021) despite some early work on this topic (Schuster and Schröder 1990a; Schuster and Schröder 1990b).

The objective of this study was to assess in a field experiment the impact of three individual PPPs containing each one a.i. (the herbicide clopyralid, the insecticide zeta-cypermethrin, and the fungicide pyraclostrobin) applied once or sequentially at two different concentrations on soil microbiota and nematode endpoints.

## MATERIALS AND METHODS

### Study site and pesticide application

The field study took place in an agricultural area in Flörsheim am Main (50° 00’ 58.1“ N 8° 23’ 41.0“ E), Germany. The soil is silty loam, characterized by 20.6% of clay, 24.8% of silt and 54.6% of sand. Before the beginning of the experiment, the soil was harrowed to clear the surface from crop residues. The field was not irrigated, and weeds were not removed during the experiment.

The field site was partitioned into 27 plots, each measuring 2.5 m by 2.5 m with an interplot distance of 2 m. Each plot was treated with either a single a.i. once, the sequential application of all three a.i., or no application (control). Plots were assigned to treatments following a Latin square design (Fig. S1). The a.i were applied as commercial formulations at two different application rates (1x or 10x of the agronomical dose of the respective product). Applications were performed with a hydraulic sprayer (Agrotop GmbH, Obertraubling, DE) with a water volume of 350 L/ha. The studied a.i were clopyralid (CAS 1702-17-6, herbicide, commercial formulation: Cliophar, a.i concentration: 100 g/L, agronomical dose: 1.5 L/ha), zeta-cypermethrin (CAS 1315501-18-8, insecticide, commercial formulation: Minuet® 10EW, a.i concentration: 100 g/L, agronomical dose 0.75 L/ha) and pyraclostrobin (CAS 175013-18-0, fungicide, commercial formulation: Comet® 200, a.i concentration: 200 g/L, agronomical dose: 1.25 L/ha).

The application and sampling scheme is illustrated in Figure 1. The single as well as the sequential applications were performed at one-week intervals, first clopyralid (day 1), then zeta-cypermethrin (day 8) and last pyraclostrobin (day16). All plots were sampled before the first application (day 0). All control and mixture treatment plots were sampled at all sampling days over the duration of the experiment. The plots that received a single application of one a.i. were sampled one day before and 7 as well as 28 days after the respective applications. Analytical samples were taken by randomly placing one cellulose filter paper within a Petri dish (9 cm diameter) on two plots per each dose during PPPs application. The filter papers were pooled and stored at −20 °C until analytical analysis at TZW (DVGW-Technologiezentrum Wasser, Karlsruhe, Germany). The measured a.i. concentrations are expressed as % of the nominal concentration.

**Figure 1.**
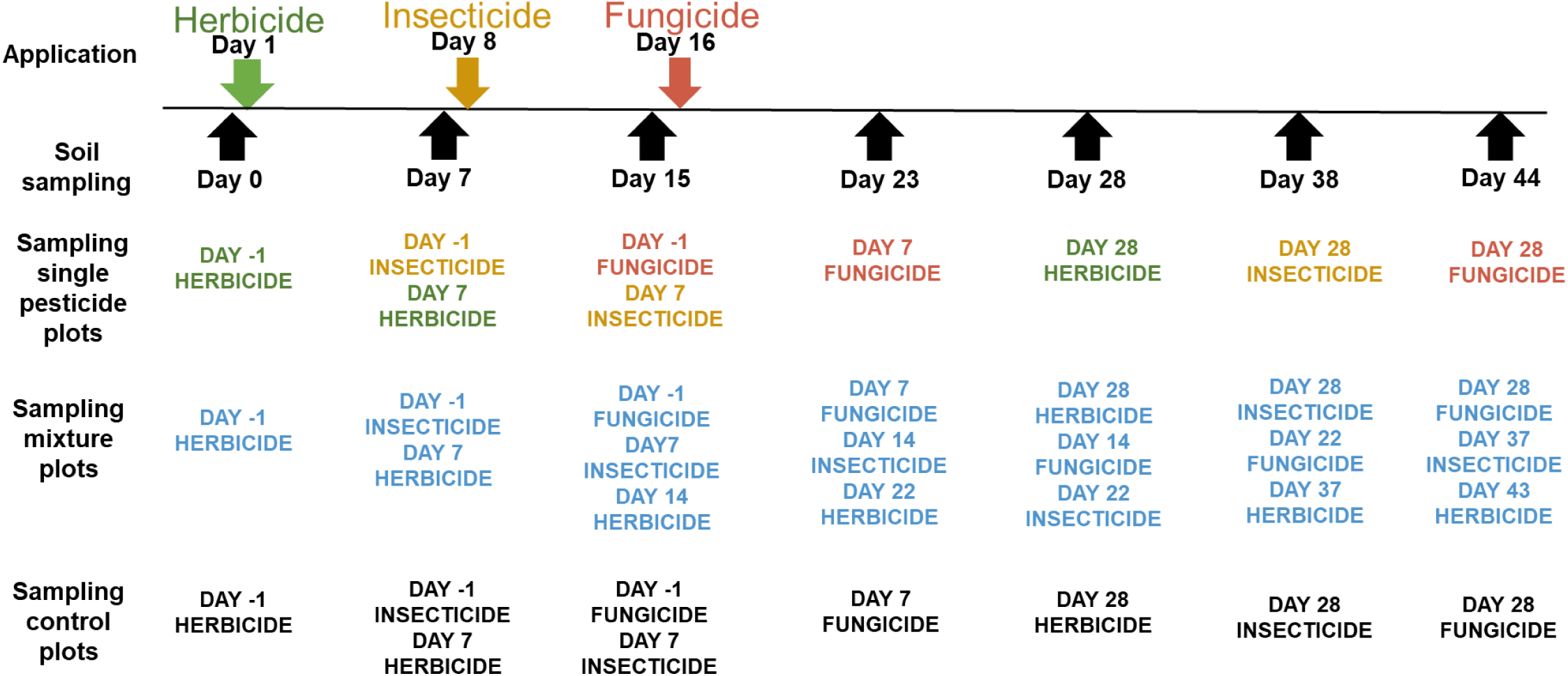
Schematic representation of the treatment and sampling schedule. Single pesticide treatment plots sampled one day before the respective pesticide applications and day 7 and 28 after the respective applications. Control and mixture plots sampled at each timepoint over the 44 days

At each sampling, five soil cores of 2 cm diameter and 5 cm depth were pooled per plot to obtain a representative composite sample. The soil was then sieved to 4 mm and portioned. Subsamples for nematode analysis were stored at 4°C for up to ca. 4 weeks, and subsamples for DNA extraction were stored at −20°C for several months. Subsamples of each plot taken on day 0 were analyzed for pH, organic material (OM), organic carbon (OC), cation exchange capacity (CEC), C/N ratio (RATIOCN), and total nitrogen (NTOT) content.

### Nematode extraction and counting

Nematodes were then extracted from 100 g fresh soil using a modification of Cobb’s decanting and sieving method (1918). Briefly, 100 g of fresh soil were mixed with 800 mL of water and soaked for about 30 minutes to detach the nematodes from soil particles. Thereafter, the soil suspension was stirred and decanted for 30 seconds. The supernatant was then poured in a collection plastic bowl. This procedure was repeated 3 times. At the end the remaining sediment was discarded.

The supernatant with nematodes went through several filters with decreasing pore size: 1000, 350, 175, 100 and 45 µm (repeated 4 times). The collected impurities on the 1000 µm filter were rinsed and discarded. The nematodes on the 350, 175, 100 and 45 sieves were rinsed with tap water and transferred to a separate bowl. The nematode suspension was then transferred to an extraction sieve with 3 layers of milk filters. The nematodes suspension was incubated for 46 hours at room temperature to allow the migration of the nematodes through the filters to the collection tray. Nematodes were counted from three 5 mL aliquots and expressed as nematodes number/100 g of soil dry weight. DNA extraction

The DNA was extracted from 250 mg of dry weight soil using the DNeasy PowerSoil DNA extraction Kit (Qiagen) following the kit’s instructions. The DNA was quantified with the Quant-iT™ dsDNA Assay Kit (Thermo Fischer Scientific, France), and stored at −20°C until use.

### Assessment of microbial community composition

Soil microbiota composition was assessed by considering both bacterial and fungal communities’ compositions and diversities monitored via amplicons Illumina Miseq 2×250 bp. 16S rRNA V3-V4 hypervariable region and the Internal Transcribed Spacer 2 (ITS2) region of the ribosomal RNA gene cluster, were targeted for bacterial and fungal communities, respectively. For both target (341F (5’-CCTACGGGRSGCAGCAG-3’) and 805R (5’-GACTACCAGGGTATCTAAT-3’) for bacteria and ITS3F (5’-GCATCGATGAAGAACGCAGC-3’) and ITS4R (5’-TCCTCSSCTTATTGATATGC-3’) for fungi) a two-steps PCR amplification method was applied (Berry et al. 2011). In the first step of both bacterial and fungal communities were amplified with following the thermal cycling condition: 3 min at 98°C, followed by 25 cycles ×(15 sec at 95°C, 30 sec at 55°C and 30 sec at 72°C), and 10 min at 72°C for bacteria and 3 min at 95°C, followed by 30 cycles ×(15 sec at 95°C, 30 sec at 55°C and 30 sec at 72°C), and 7 min at 72°C for fungi. In a second step, the amplicons duplicates of the first bacterial and fungal PCR were pooled and used as a template to link a unique combination of multiplex primer pair to the overhang adapters (forward: TCGTCGGCAGCGTCAGATGTGTATAAGAGACAG, adapter: GTCTCGTGGGCTCGGAGATGTGTATAAGAGACAG) following thermal cycling conditions 3 min at 98°C, then 8 cycles ×(15 sec at 98°C, 30 sec at 55°C, 30 sec at 72°C), and 10 min at 72°C.

The bacterial and fungal amplicons from the second PCR were visualized in a 2% agarose gel for size and amplification check. They were then pooled together and purified with the SequalPrep Normalization Plate kit (Invitrogen, Frederick, MD, USA). Sequencing was performed by GenoScreen (Lille, FR) on MiSeq (Illumina 2×250 bp) using the MiSeq reagent kit v2 (500 cycles).

### Bioinformatics analysis

The sequence data were analyzed using an in house developed Jupyter Notebooks (Kluyver et al. 2016) piping together different bioinformatics tools. Briefly, for both 16S rRNA and ITS data, R1 and R2 sequences were assembled using PEAR (Zhang et al. 2014) with default settings. Further quality checks were conducted using the QIIME pipeline (Caporaso et al. 2010) and short sequences were removed (< 400 bp for 16S rRNA and for ITS). Reference-based and de novo chimera detection, as well as clustering in OTUs were performed using VSEARCH (Rognes et al. 2016) and the adequate reference databases (SILVA’ representative set of sequences for 16S rRNA and SILVA’s ITS2 reference dynamic dataset for ITS). The identity thresholds were set at 94% for 16S data based on replicate sequencing of a bacterial mock community containing 40 bacterial species, and 97% for ITS data for which we did not have a mock community. For 16S rRNA data, representative sequences for each OTU were aligned using MAFFT (Katoh and Standley 2013) and a 16S phylogenetic tree was constructed using FastTree (Price et al. 2009). Taxonomy was assigned using BLAST (Altschul et al. 1990) and the SILVA reference database v138 for 16S rRNA (Quast et al. 2013). For ITS, the taxonomy assignment was performed using BLAST (Altschul et al. 1990) and the UNITE reference database (Abarenkov et al. 2021). Diversity metrics, that is, Faith’s Phylogenetic Diversity (PD) for 16S rRNA data only (Faith 1992), richness (observed species-OS) and evenness (Simpson’s reciprocal index-SI), describing the within-sample microbial communities’ structure (α-diversity) were calculated based on rarefied OTU tables (22000 and 15000 sequences per sample for 16S rRNA and ITS respectively). Bray-Curtis and UniFrac distance matrices (Lozupone and Knight 2005) were also computed to detect global variations in the composition of microbial communities between samples (β-diversity) for ITS and 16S rRNA, respectively.

### Quantification of microbial communities

The total abundances of bacteria (16S rRNA), fungi (ITS), ammonia oxidizing archaea (AOA) and bacteria (AOB), and comammox A (COMAA) and B (COMAB) were quantified through real-time quantitative PCR (qPCR) using the ViiA7TM thermocycler (Life Technologies, Carlsbad, CA, USA). The reaction volume was 15 µL per sample and contained 7.5 µL of Takyon Low Rox SYBR MasterMix dTTP blue (Eurogentec, Seraing, Belgium), 1 µM of each primer, 250 ng of T4 gene 32 protein (MP Biomedicals, Illkirch, France), and 1.5 ng of DNA. Total bacterial and fungal community were quantified according to Muyzer et al. (1993) and White et al. (1990), respectively. The abundance of AOA and AOB were estimated to target the amoA gene (Bru et al. 2011). COMAA and COMAB were quantified targeting the clade A and B amoA (Pjevac et al. 2017). Non-template controls (NTC) were analyzed too, and no quantification could be detected. Standard curves were created with serial dilutions of linearized plasmids containing cloned genes from bacterial strains or environmental clones. Two independent runs were performed on each gene quantification. The qPCR’s efficacy ranged from 78% to 107%. The final abundances were expressed in number of numbers of copies/g of soil dry weight. Prior to analysis, an inhibition test was performed mixing the soil DNA with a control plasmid (pGEM-T Easy Vector, Promega, Madison, WI, USA) or water to detect the presence of inhibitors of the qPCR assays. No inhibition could be detected.

### Statistical analysis

Statistical analyses were performed using the software R studio version 2023.12.1+402. We first assessed the spatial variability of soil properties within the experimental field (i. e. pH, OM, OC, NTOT, CEC and RATIOCN), employing ANOVAS with the plot positions (X and Y coordinates) as explaining factors.

To assess the impact of single and mixture treatments on soil microbial communities and nematode abundance, we used multiple generalized linear mixed models (GLMMs) using the glmmTMB function from the glmmTMB package (version 1.1.8), which accounts for both fixed effects and random effects. Since we observed a spatial variability for certain soil properties, we incorporated them as fixed effects across all models. This approach enables us to distinguish the treatment effect from those of the soil properties spatial variability. To reduce collinearity among variables, only soil properties with low correlation were included in the models (i. e. OM, pH, RATIOCN and CEC) (Fig. S2). To account for the temporal variability, we included the sampling day into the models as random effects. Three models were then constructed to evaluate the effects of i) pesticides mixture across all sampling days, ii) single treatments (i. e. for herbicide, insecticide and fungicide altogether), and iii) single and mixture treatments (i. e. for herbicide, insecticide and fungicide, individually). For models ii) and iii), we only considered the day preceding the application of the respective pesticide (day −1), 7 and 28 days following the pesticide application (day 7, day 28, respectively). Normality and homogeneity of the residual distribution were verified, and log-transformations were performed when necessary for gene copy abundances. We then implemented multiple pairwise comparisons for each significant treatment-dose effect using the emmeans function from the emmeans package (version 1.10.0). The p-values were then adjusted using the false discovery rate (FDR) method (Benjamini and Hochberg 1995).

We then conducted a Partial Redundancy Analysis (P-RDA) model to explore the impact of the Treatment-by-Dose and soil properties (explanatory variables) on the composition of the bacterial and fungal communities (response variables), while considering the variation due to the sampling day as a covariate. To perform this analysis, low-abundance OTUs were filtered out, retaining only those representing > 0.01% of the sequences across all samples which resulted in 1280 OTUs for 16S and 587 OTUs for ITS. The significance of the P-RDA model and its variables was assessed using the anova.cca function (with 1000 permutations) in the vegan package.

Additionally, the nematode abundance data were further analyzed as described by Fernandes et al. (submitted) for microarthropod data. Briefly, the post-application nematode abundances at day 7 and day 28 were divided by the initial abundance (day −1) to calculate the short-term and long-term ratio of abundance changes, respectively. The abundance ratios were log-transformed and statistically analyzed with multiple one-way-ANOVAs. Each statistical analysis was carried out on the single treatments (i. e. for herbicide, insecticide, fungicide and mixture, individually) on either the short-term or long-term ratio. Additionally, the mixture treatment was analyzed with a two-ways-ANOVA, using treatment-dose and the ratio (short-term and long term) as factors. This analysis enables us to compare the changes in nematode to that of the microarthropods explored by Fernandes et al. (submitted), exploring the tested PPPs effects on multiple layers of the soil food web.

## RESULTS

### Spatial variability of plot properties and verification of application rates

All measured soil properties, except for the pH, were significantly correlated with their X and Y spatial plot coordinates (p ≤ 0.05), indicating heterogeneity in OC, OM, NTOT, RATIOCN and CEC across the field site (Tab. 1 and Fig. S3). The OC, OM, and NTOT were significantly influenced by both X and Y coordinates, while RATIOCN and CEC varied only along the Y and X axis, respectively.

**Table 1.** Result of the ANOVA analysis of the spatial variability of soil physicochemical properties (OM, organic matter; pH, soil pH; RATIOCN, ratio C/N; CEC, cationic exchange capacity) in relation to plot coordinates

| PLOT | pH |  |  | OC |  |  | OM |  |  |
| --- | --- | --- | --- | --- | --- | --- | --- | --- | --- |
| COORDINATE | Df | F | *p* | Df | F | *p* | Df | F | *p* |
| X<sup>a</sup> | 2 | 0.936 | 0.413 | 2 | 4.095 | <0.05 | 2 | 4.374 | <0.05 |
| Y<sup>b</sup> | 8 | 1.021 | 0.459 | 8 | 6.435 | <0.05 | 8 | 6.484 | <0.05 |

| PLOT | NTOT |  |  | RATIOCN |  |  | CEC |  |  |
| --- | --- | --- | --- | --- | --- | --- | --- | --- | --- |
| COORDINATE | Df | F | *p* | Df | F | *p* | Df | F | *p* |
| X<sup>a</sup> | 2 | 5.316 | <0.05 | 2 | 0.830 | 0.454 | 2 | 11.239 | <0.05 |
| Y<sup>b</sup> | 8 | 2.960 | <0.05 | 8 | 4.673 | <0.05 | 8 | 1.396 | 0.271 |

<sup>a</sup> coordinate X
<sup>b</sup> coordinate Y

The analytically determined amounts of the a.i. in the paper filters were related to the nominally applied amount and reached 28.0% and 26.7% for clopyralid-1× and clopyralid-

10×, respectively, 68.1% and 92.6% for cypermethrin-1× and cypermethrin-10×, respectively, and 74.8% and 69.1% for pyraclostrobin-1× and pyraclostrobin-10×, respectively.

### Effects on bacterial and fungal community structure

Results from the five GLMM models used to evaluate the effects of single or sequential applications of PPPs on bacterial α-diversity showed no PPPs impact (Tab. S1, Figs. S4 & S5 & S6). Regarding fungal α-diversity (Tab. S2), the SI was the most sensitive endpoint (Fig. 2). Even though fluctuating overtime, the pairwise comparisons showed significantly higher values in the Mixture-1× treatment compared to the Control. Pairwise comparisons of the single application model (Fig. 2-panel B1) showed significantly lower SI in Control samples compared to the Herbicide-10×. When comparing the herbicide single application to the mixture application, fungal SI showed consistently lower values in Control compared to Herbicide-1× and Mixture-10×. No significant differences could be detected for OB (Fig. S7).

**Figure 2.**
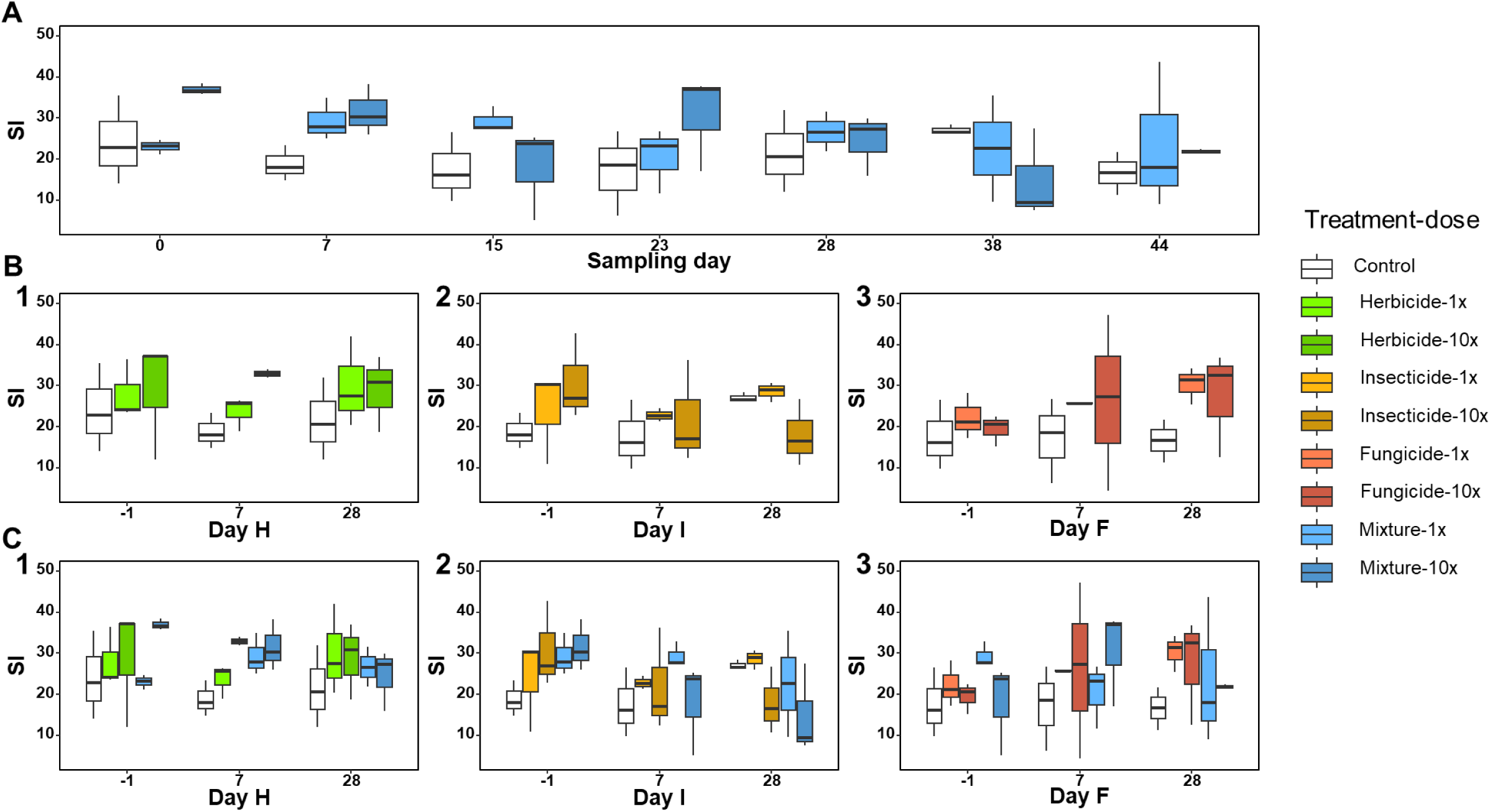
Fungal Simpson index (SI) for the mixture (panel A), the single pesticides (panel B, 1-2-3) and the mixture and single treatments together (panel C, 1-2-3) at the different sampling dates. Day H, Day I and Day F represent the sampling day relative to the individual pesticide application (−1: 1 day before respective application; 7: 7 days after respective application; 28: 28 days after respective application). The treatments are control, herbicide (clopyralid), insecticide (zeta-cypermethrin), fungicide (pyraclostrobin) and sequential mixture (mixture), all tested at 1× or 10× the agronomical dose. Shown are the medians, the lower and upper hinges which correspond to the first and third quartiles

The partial-RDA analysis (Fig. 3) showed that soil physio-chemical properties and PPPs treatments explained 21% and 18% of the observed constrained variance in the bacterial and fungal communities composition, respectively (Tab. 2). The anova.cca function shows that 4% and 20% of the variance was inferred to the PPPs for bacterial and fungal community composition, respectively (Tab. 3). These two observations are visually exemplified in the fungal community partial-RDA plot (Fig. 3-panel B) where the Control plots tend to cluster on the opposite of most of the treated plots, while Control plots did not cluster as much for bacterial community composition (Fig. 3-panel A). For both bacterial and fungal community composition the length of the arrows in the partial-RDA analysis indicates that the influence of soil properties (particularly pH, CEC, OM, and RATIOCN) is much stronger than the influence of PPP application.

**Figure 3.**
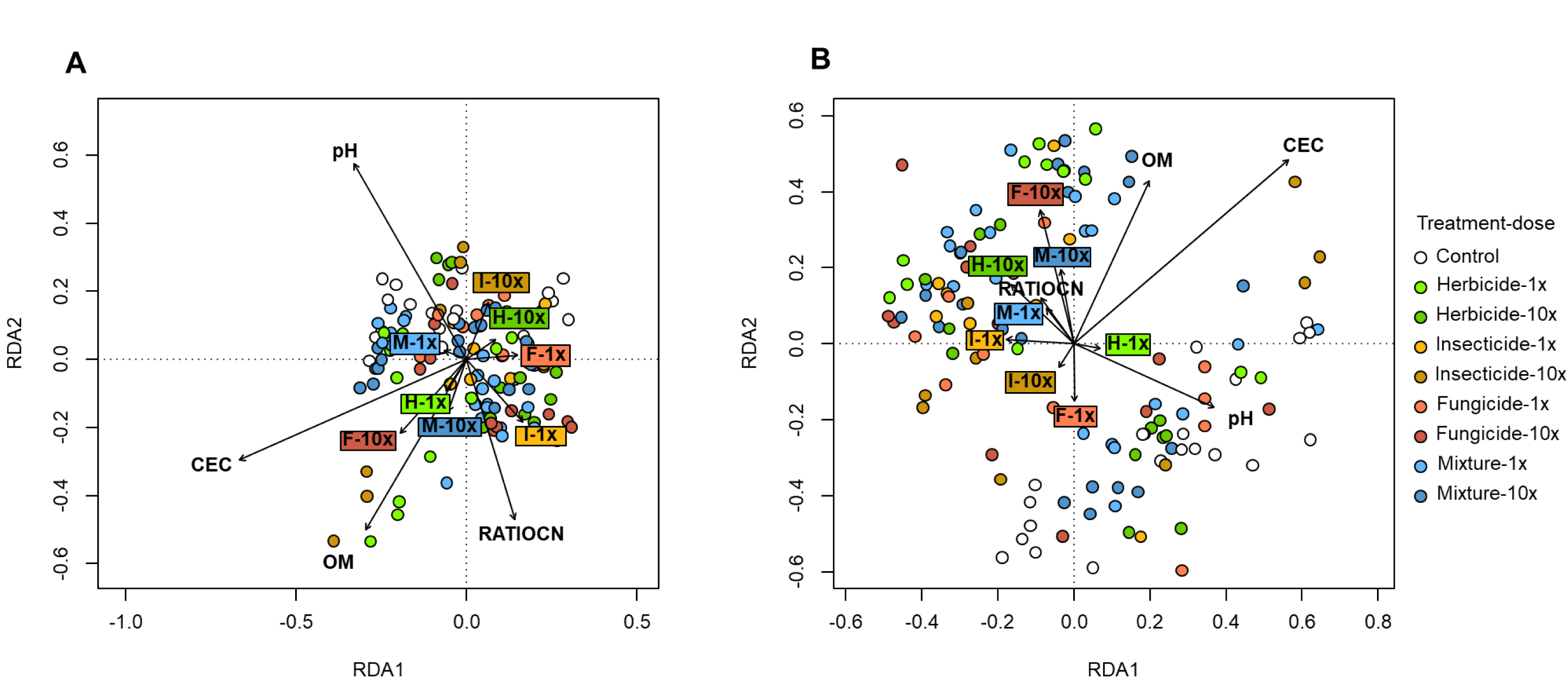
Partial Redundancy Analysis (P-RDA) of the bacterial (panel A, 16S rRNA amplicon) and fungal (panel B, ITS amplicon) community composition in dependence of pesticide treatments (control, herbicide (clopyralid), insecticide (zeta-cypermethrin), fungicide (pyraclostrobin) and their sequential mixture (all at 1x or 10x the agronomical dose)) as well as soil properties. The different colors represent the different treatment-dose combinations. The length and direction of the arrows illustrate the contribution of the factors to the observed variation between the communities

**Table 2.** Results of the conditioned, constrained and unconstrained variances explained by the P-RDA model for bacterial and fungal communities’ composition analyses assessed by 16S rRNA and ITS metabarcoding, respectively

|  | Bacteria (16S rRNA) |  | Fungi (ITS) |  |
| --- | --- | --- | --- | --- |
| Variance | Inertia | Proportion | Inertia | Proportion |
| Total | 0.05 | 1.00 | 0.21 | 1.00 |
| Conditioned | 0.01 | 0.08 | 0.01 | 0.05 |
| Constrained | 0.01 | 0.21 | 0.04 | 0.18 |
| Unconstrained | 0.04 | 0.70 | 0.16 | 0.77 |

**Table 3.**
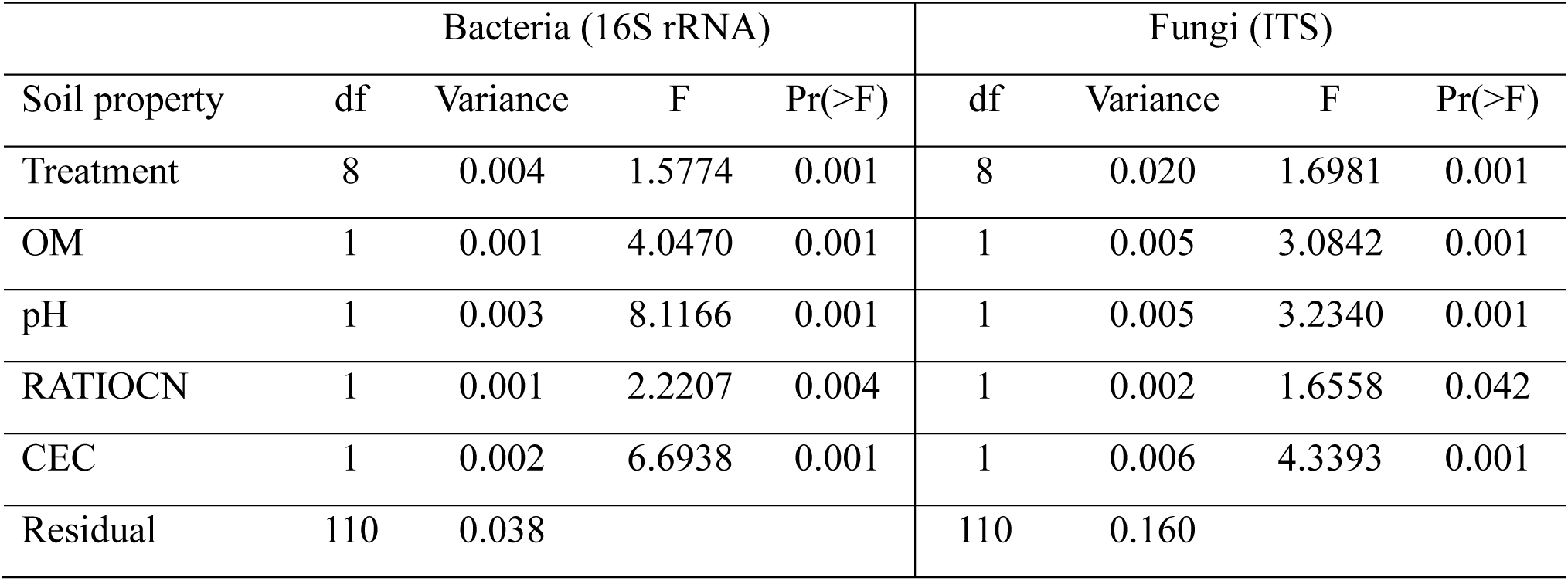
Results of the anova.cca analysis on the constrained variance for the soil physicochemical parameters (OM, organic matter; pH, soil pH; RATIOCN, ratio C/N; CEC, cationic exchange capacity) bacterial and fungal communities’ composition analyses assessed by 16S rRNA and ITS metabarcoding, respectively

### Effects on the abundance of bacteria, fungi and nitrifiers

The 16S and ITS abundances displayed overall a relatively small range of variations (from 1.4×10^8^ copies number/g dry soil to 1.2×10^9^ copies number/g dry soil for 16S, and from 9.0×10^7^ copies number/g dry soil to 7.4×10^8^ copies number/g dry soil for ITS, Tab. S3, Figs. S8 & S9). Significant differences were detected when comparing the fungicide and mixture treatments. Specifically, Mixture-1× displayed lower 16S abundance compared to the Control and Fungicide-10×. Significantly lower ITS abundance was detected in the insecticide vs mixture analysis, where the Mixture-10× had a significantly higher ITS abundance compared to Control, Insecticide-1× and Insecticide-10×.

The AOA/AOB×100 remained stable in the Control treatment over the duration of the experiment, while it sharply increased at day 15 and 44 for the Mixture-1× treatment, resulting in an overall significant higher AOA abundance compared to Mixture-10× (Tab. S3, Fig. S10-panel A). The AOA/AOB×100 quantified in the Mixture-1× was also significantly higher compared to Fungicide-10× and Mixture-10× according to the fungicide vs mixture analysis.

After 28 and 38 days the AOB/16S×100 measured in Mixture-10× treatment were similar to that observed on day 0, thus indicating its recovery at the end of the experiment (Tab. S2, Fig. 4-panel A). However, the AOB/16S×100 measured in Mixture-1× treatment was statistically lower than that of Mixture-10× treatment all over the duration of the experiment, and also significantly lower compared to Fungicide-10× in the fungicide vs mixture comparison.

**Figure 4.**
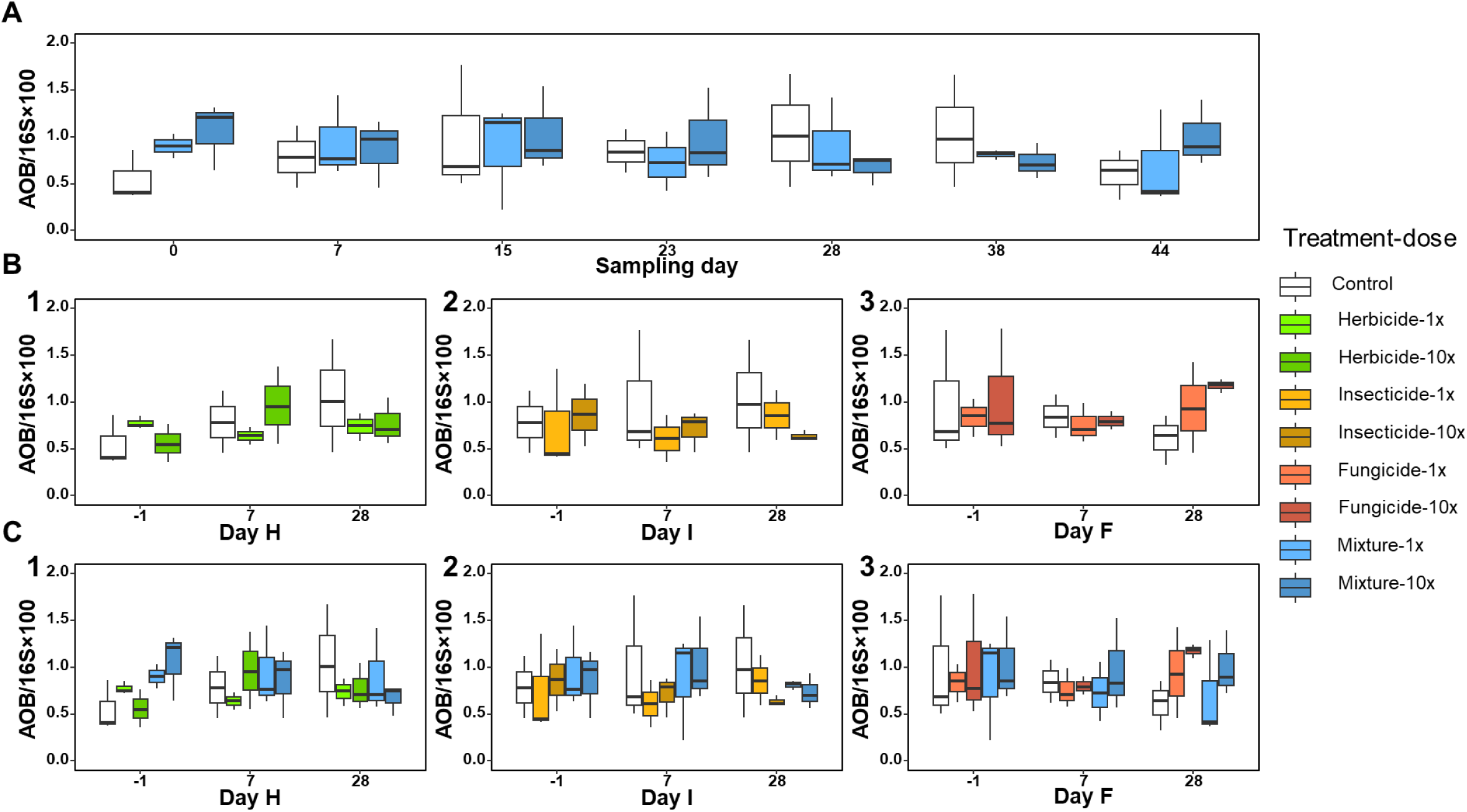
Relative abundance of ammonia oxidizing bacteria (AOB/16S*100) for the mixture (panel A), the single pesticides (panel B, 1-2-3) and the mixture and single treatments together (panel C, 1-2-3) at the different sampling dates. Day H, Day I and Day F represent the sampling day relative to the individual pesticide application (−1: 1 day before respective application; 7: 7 days after respective application; 28: 28 days after respective application). The treatments are control, herbicide (clopyralid), insecticide (zeta-cypermethrin), fungicide (pyraclostrobin) and sequential mixture (mixture), all tested at 1× or 10× the agronomical dose. Shown are the medians, the lower and upper hinges which correspond to the first and third quartiles

The COMAA/16S×100 (Tab. S3, Fig. S11-panel A) and COMAB/16S×100 (Tab. S3, Fig. S12-panel A) exhibited similar patterns characterized by a sharp increase in Mixture-1× on day 15. This increase led to a significantly higher COMAB abundance in Mixture-1× compared to the Control. Both COMAA/16S×100 and COMAB/16S×100 was significantly increased by the Herbicide-1× treatment as compared to the Control. Interestingly, in the comparison between the herbicide and the mixture, a different Herbicide-1× effect was observed on the two comammox communities. For COMAA it led to a significantly lower COMAA compared to the Control, Herbicide-10×, and Mixture 1×. In contrast, in COMAB Herbicide-1× caused a significant enhancement compared to the Control.

Overall, the abundance of bacteria, fungi and nitrifiers indicated limited temporal variations over the duration of the experiment, with idiosyncratic impacts of pesticide applications of relatively low effect sizes that might not be biologically relevant.

### Effects on nematode abundance

The nematode abundance (Fig. 5) remained stable in control until day 23, when it increased and stabilized at a higher level for the remaining duration of the experiment (Tab. S4, Fig. 5-panel A). Results of the GLMM model of the mixture analysis show averaged over time significantly higher nematode abundance in the Control and Mixture-10× compared to Mixture-1×. In the insecticide vs mixture comparison, Mixture-10× had a significant higher nematode abundance compared to Insecticide-1×, Insecticide-10× and Mixture-1×. In the fungicide vs mixture, Fungicide-10× at day 7 has a sharp increase in nematode abundance, which decreased on day 28, and resulted in significantly higher nematode abundance compared to Fungicide-1× and Mixture-1×. The increase in the nematode abundance observed in the Fungicide-10× samples at day 7 was not observed anymore at day 28 and consequently nematode abundance in Fungicide-10× was not anymore different from those of Fugicide-1× and Mixture-1× samples.

**Figure 5.**
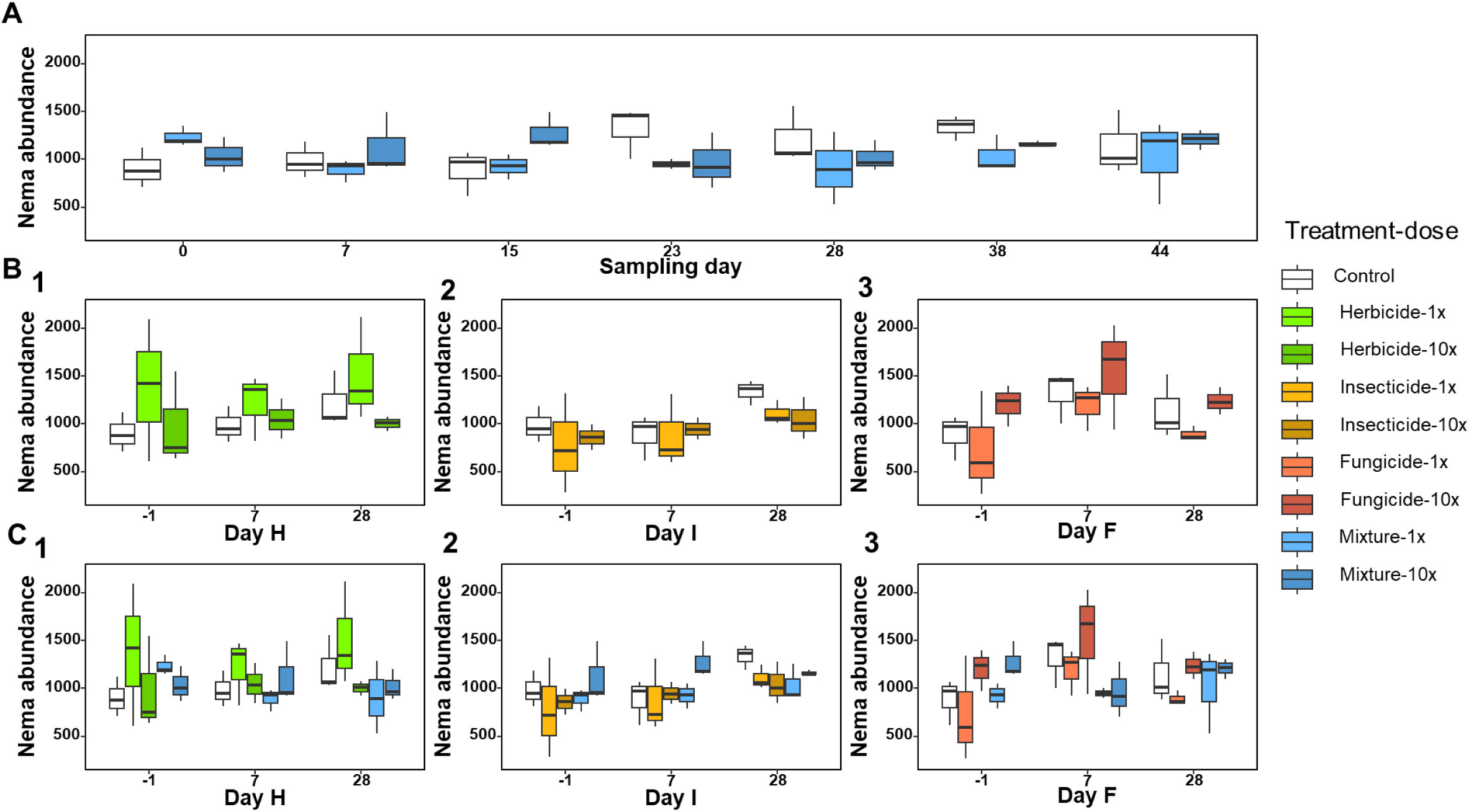
Nematode abundance for the mixture (panel A), the single pesticides (panel B, 1-2-3) and the mixture and single treatments together (panel C, 1-2-3) at different sampling dates. Day H, Day I and Day F represent the sampling day relative to the individual pesticide application (−1: 1 day before respective application; 7: 7 days after respective application; 28: 28 days after respective application). The treatments are control, herbicide (clopyralid), insecticide (zeta-cypermethrin), fungicide (pyraclostrobin) and sequential mixture (mixture), all tested at 1× or 10× the agronomical dose. Shown are the medians, the lower and upper hinges which correspond to the first and third quartiles

The analysis of the nematode abundance changes on the short-term and longer-term duration did not indicate effects different from the above analysis.

## DISCUSSION

The aim of this study was to investigate the effect of the sequential application of three PPPs on a bare agricultural field, focusing on the structure and diversity of the bacterial and fungal soil communities, as well as on the total abundances of bacteria, fungi and nematodes. We employed a combination of molecular tools for bacterial and fungal community analyses, and microscopic counting measuring the nematode abundances. We also assessed the spatial heterogeneity of several soil physio-chemical properties across the field to better understand their potential influence on the observed outcomes.

Overall, we observed a large spatial heterogeneity of soil physio-chemical properties that explained a large part of the relatively high variability between biological triplicates for many measured endpoints. Belowground communities are indeed deeply influenced and shaped by soil physio-chemical characteristics, which can be variable in a field site even at relatively small scale (Goovaerts 1998). We therefore included plot soil properties into the statistical analyses, to properly evaluate the effects of the PPPs treatments on the measured endpoints. Organic matter was the most variable property within the field site, despite mechanical pre-treatment. Decomposition of organic matter is directly mediated by microbes (Liang et al. 2019) and organic matter content influences the abundance of specific microbial groups and microbial community structure in soil (Inagaki et al. 2023, Yao et al. 2024). Hence, within-field heterogeneity of organic matter drove the variability of microbial communities among plots. While subsampling of plots and implementing the Latin square design in this experiment to decrease the bias resulting from the correlation of measured biological endpoints with soil properties, the statistical power of the experiment was still impacted by the high variability.

Accordingly, results of the partial-RDA of the bacterial community structure showed that most of the constrained variance was explained by variation in the soil physio-chemical properties, rather than by PPPs application. Many studies indicate soil pH as the main driver of bacterial community composition (Lauber et al. 2008, 2009; Shen et al. 2013; Philippot et al. 2023), as it influences the conditions that enable the microbes to survive (Jin and Kirk 2018). In our study, soil pH was homogenous within the field, while organic matter content and cation exchange capacity appeared to be equally stronger drivers.

Still, fungal abundance and community composition were the most responsive to PPPs application among the measured biological endpoints in this study. Both sequential mixture and herbicide treatments led to significant increases in the fungal abundance and modifications of the fungal diversity as compared to the Control. The ecotoxicological impact of pesticides on fungal communities is less studied than that of bacteria. Generally, fungi harbor vast enzymatic arsenals able to degrade various organic compounds which make them resilient to PPPs application (Kalia and Gosal 2011). In agreement with our findings, several recent studies reported adverse PPPs effects on the fungal community diversity and composition (Streletskii et al. 2022, 2023), that could be associated with undirect effects on obligate symbiotic species i.e. arbuscular mycorrhiza. Karpouzas et al. (2014) described how the direct effect of herbicide on plant caused an indirect reduction of mycorrhizal colonization and richness due to missing symbiotic interaction. The variable responses to pesticide applications reported in the literature could be influenced by the community’s historical exposure to pesticides, potentially indicating existing adaptation to certain a.i. entering in the composition of PPPs.

The N cycle is known to be sensitive to pesticide as the nitrification guild due to the taxonomic narrowness of ammonia oxidizers (Karas et al. 2018; Karpouzas et al. 2022; Sim et al. 2022). In our study, the abundance of comammox increased in response to pesticide application, especially in the case of COMAB which exhibits sensitivity to the herbicide already at low dose. This finding is in contrast with other studies that reported adverse effects of herbicides on the N cycle (Damin and Trivelin 2011).

Soil nematodes live in the water layer of the soil pores having diameter of 25-100 µm (Neher 2010). Like for soil microbes, their field distribution is deeply influenced by soil properties, among which organic carbon and pH seem to be the most impacting, together with the soil porosity (Liu et al. 2019). The substantial variation in nematode abundance in our study may stem from the pronounced variability observed in soil characteristics. Given the high variability in bacterial abundance, it is plausible that bacterivore nematodes, the most abundant feeding group in the nematode community (van den Hoogen et al. 2020), were distributed in accordance with their food sources. Even though field studies on the effects of pesticides on the nematode community are rare (Waldo et al. 2019), most of the pesticide effects are visible on the community composition rather than on the abundance. Therefore, a deeper exploration of nematode community composition, which would enable the characterization of functional aspects, would be advantageous for a comprehensive assessment of pesticide toxicity on nematodes.

Another possible explanation for the variable effects of pesticides among treatment replications could be linked to pesticide’s physio-chemical properties which influence its adsorption potential and persistence. Indeed, among pesticides used here clopyralid has the lowest absorption ability (Kfoc: 0.26–4.1 mL/g) and a low-moderate persistence in soil with half-live (DT50) ranging from 0.16 –23.7 days. This implies a high mobility in soil mediated by water flow, and also an important bioavailability for in soil living organisms. On the other hand, zeta-cypermethrin and pyraclostrobin present high Kfoc (72405-285562 mL/g and 6000-16000 mL/g, respectively) and consequently are highly adsorbed to soil components being poorly bioavailable (DT50: 6-24 days and 33 days, respectively). The higher organic matter concentration can be correlated with a higher cation exchange capacity (CEC), as the organic matter contributes to up to 60% of the CEC (Hayashi et al. 2023). CEC influences pesticide availability in soil, i. e. herbicides are generally more effective in soil with low organic matter, and so with low CEC as molecules are less likely to be absorbed and therefore remain available for their intended purpose (Kerr et al. 2004). Generally, the literature reports many cases of absorption of pesticide molecules on organic matter (Boivin et al. 2005; Yu and Zhou, 2005; Rodriguez-Cruz et al. 2007). The uneven distribution of organic matter in our experimental field, together with the different PPPs properties, might have influenced their bioavailability and subsequently exerting different effects on the microbial and nematode communities. Nevertheless, the active ingredients in the tested formulations are low toxic towards non-target organisms, another reason why we could not observe a strong pesticide effect (EFSA 2008; EFSA et al. 2018).

In-field studies estimating the effects of PPPs mixtures on soil microbes and free-living nematodes are still rare. Here, we report that the fungal community was more susceptible than the bacterial community for effects of PPPs application. The structure and diversity of the total bacterial community were mainly driven by the soil physio-chemical properties. The abundance of the nematodes was mildly affected by the PPPs applications and the effects appeared to be transient. While higher-tier ecotoxicological studies offer greater ecological relevance compared to the standard laboratory tests, challenges arise from the interpretation of pesticide effects due to the inherent and strong dependence of bacterial, fungal and nematode communities on soil characteristics and their typically high spatial heterogeneity at the field scale. Soil spatial heterogeneity should be considered as a confounding factor when deriving conclusions on the ecotoxicological effects of pesticides on soil microorganisms. At the same time, the replication of treatments and number of sub-samples per plot are technically limited in field studies, particularly when investigating numerous mixture treatments such as in the present study.

## DATA AVAILABILITY

Raw data are available on Zenodo, an open-access research data repository. The data sets can be accessed at https://doi.org/10.5281/zenodo.14882212

## ACKNOWLEDGMENTS

The project leading to this publication, as well as Camilla Drocco’s PhD, have received funding from the European Union’s Horizon 2020 research and innovation programme under the Marie Skłodowska-Curie grant agreement No 956496

