## Supplementary file for "Effects of a Sequential Application of Plant Protection Products on Soil Microbes and Free-Living Nematodes in a Field Experiment"

| | X1 | X2 | X3 |
|---|---|---|---|
| Y1 | Control | Insecticide 10x | Insecticide 1x |
| Y2 | Herbicide 1x | Herbicide 10x | Fungicide 1x |
| Y3 | Insecticide 1x | Mixture 1x | Herbicide 10x |
| Y4 | Fungicide 1x | Mixture 10x | Insecticide 10x |
| Y5 | Herbicide 10x | Control | Fungicide 10x |
| Y6 | Insecticide 10x | Herbicide 1x | Mixture 1x |
| Y7 | Fungicide 10x | Insecticide 1x | Mixture 10x |
| Y8 | Mixture 1x | Fungicide 1x | Control |
| Y9 | Mixture 10x | Fungicide 10x | Herbicide 1x |

Figure S1.

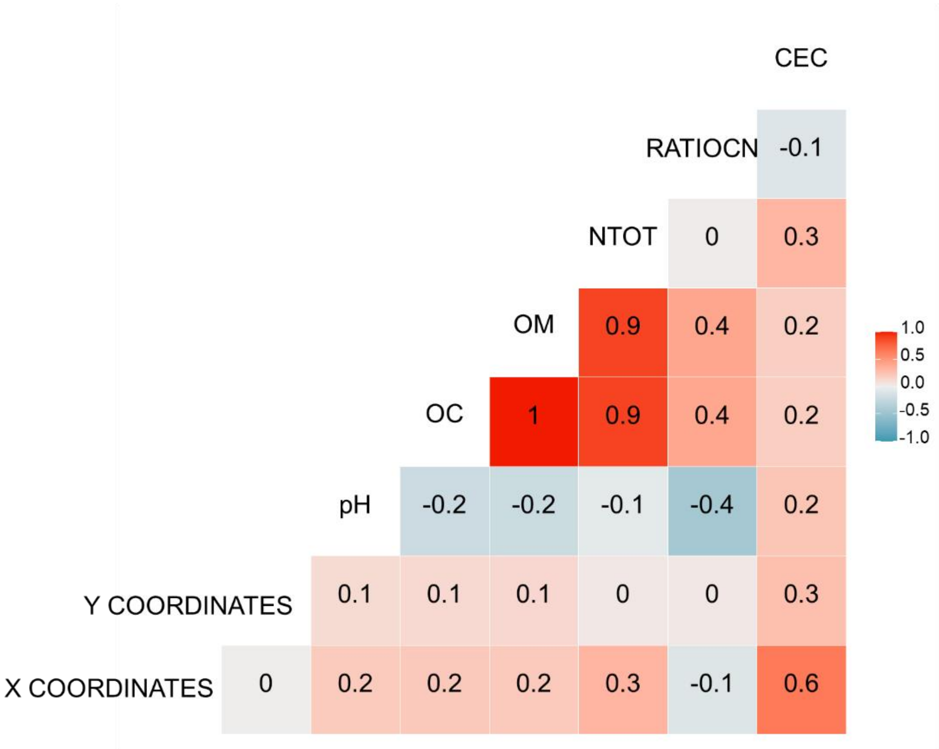
CEC
RATIOCN
NTOT
OM
OC
pH
Y COORDINATES
X COORDINATES
-0.1
0
0.3
0.9
0.4
0.2
1
0.9
0.4
0.2
-0.2
-0.2
-0.1
-0.4
0.2
0.1
0.1
0.1
0
0
0.3
0
0.2
0.2
0.2
0.3
-0.1
0.6
1.0
0.5
0.0
-0.5
-1.0

Figure S2.

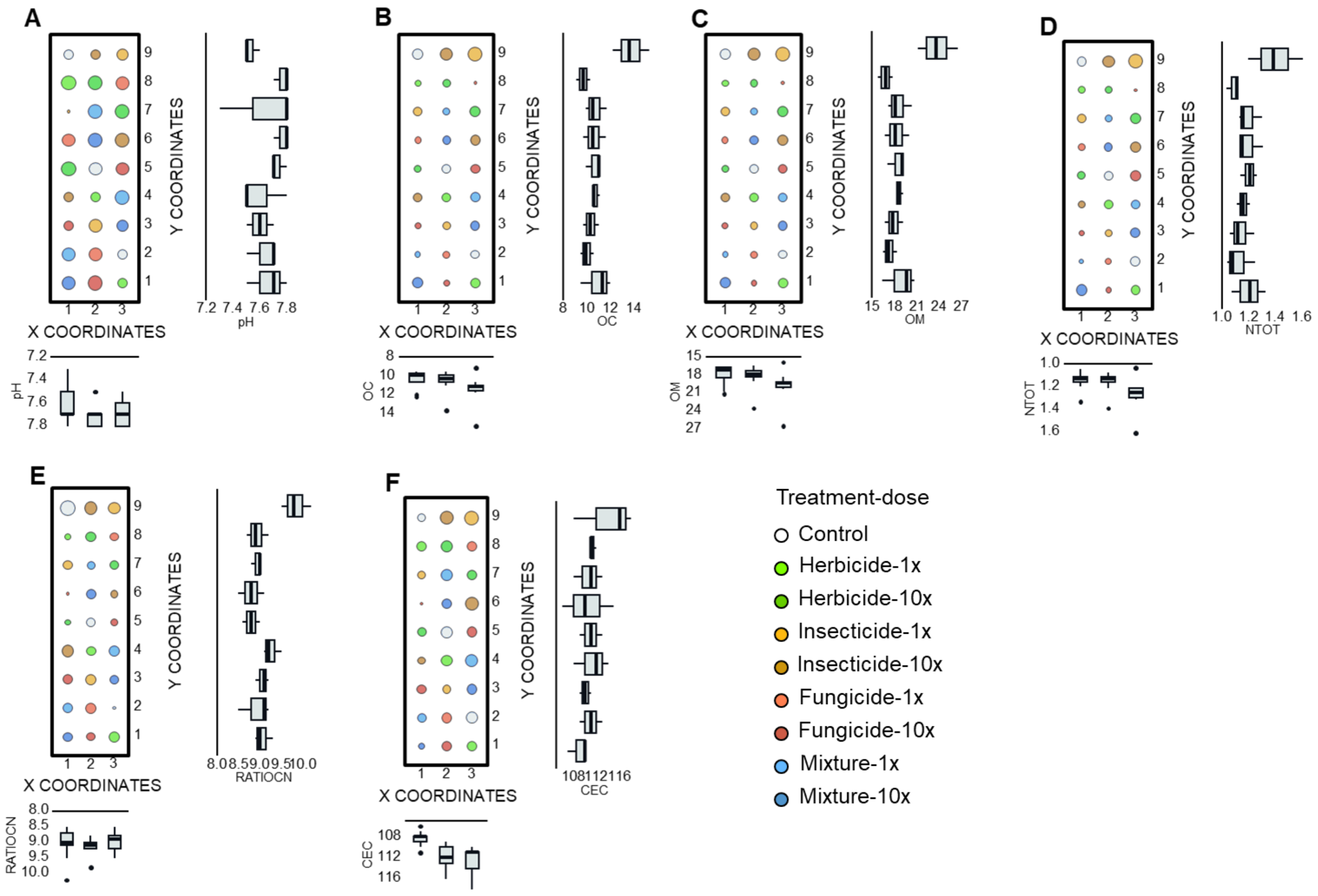

A
B
C
D
E
F
X COORDINATES
Y COORDINATES
pH
OC
OM
NTOT
RATIOCN
CEC
Treatment-dose
Control
Herbicide-1x
Herbicide-10x
Insecticide-1x
Insecticide-10x
Fungicide-1x
Fungicide-10x
Mixture-1x
Mixture-10x


Figure S3.

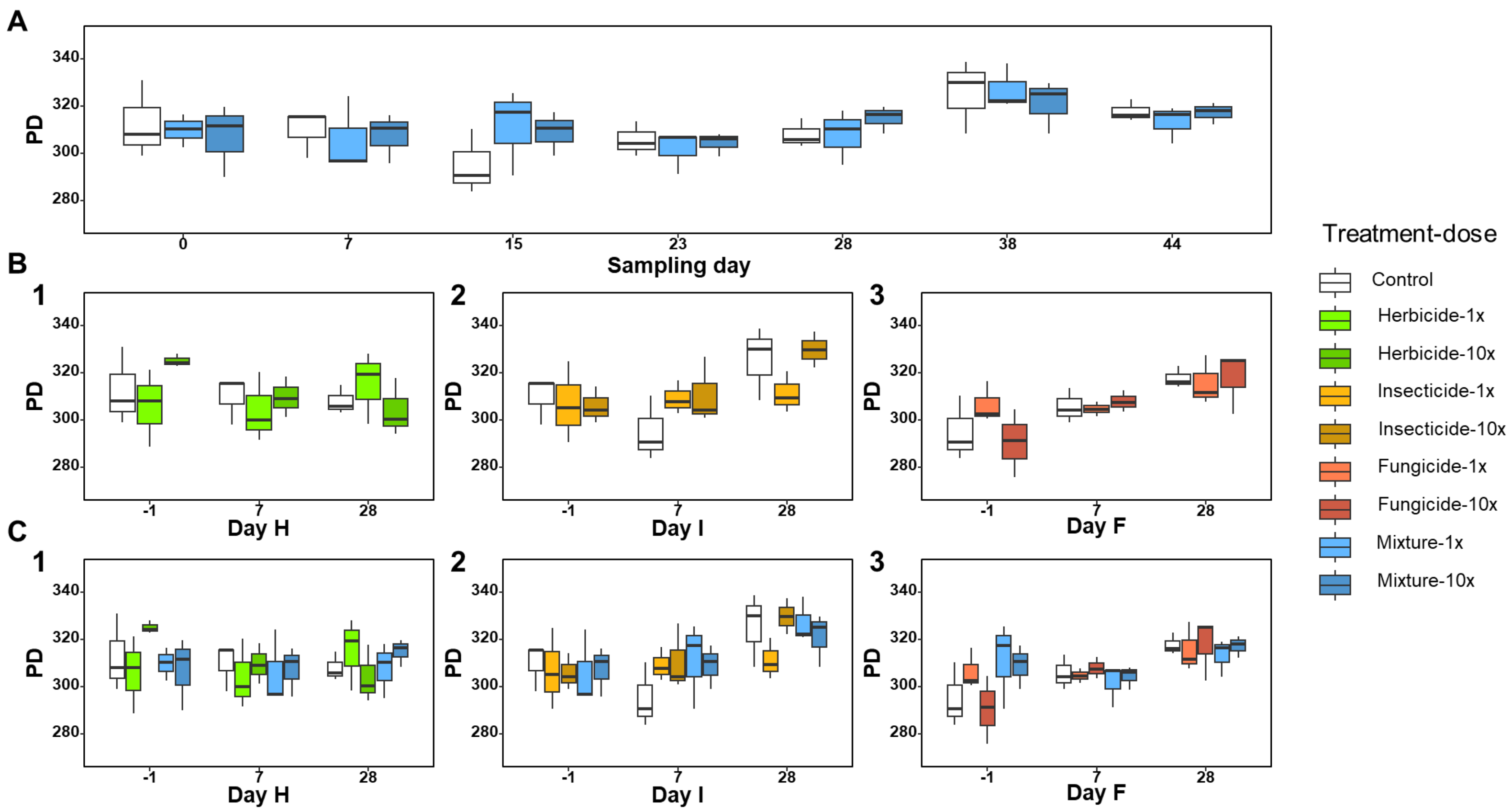
A
PD
340
320
300
280
0
7
15
23
28
38
44
Sampling day
B
1
2
3
Day H
Day I
Day F
-1
7
28
C
Treatment-dose
Control
Herbicide-1x
Herbicide-10x
Insecticide-1x
Insecticide-10x
Fungicide-1x
Fungicide-10x
Mixture-1x
Mixture-10x

Figure S4.

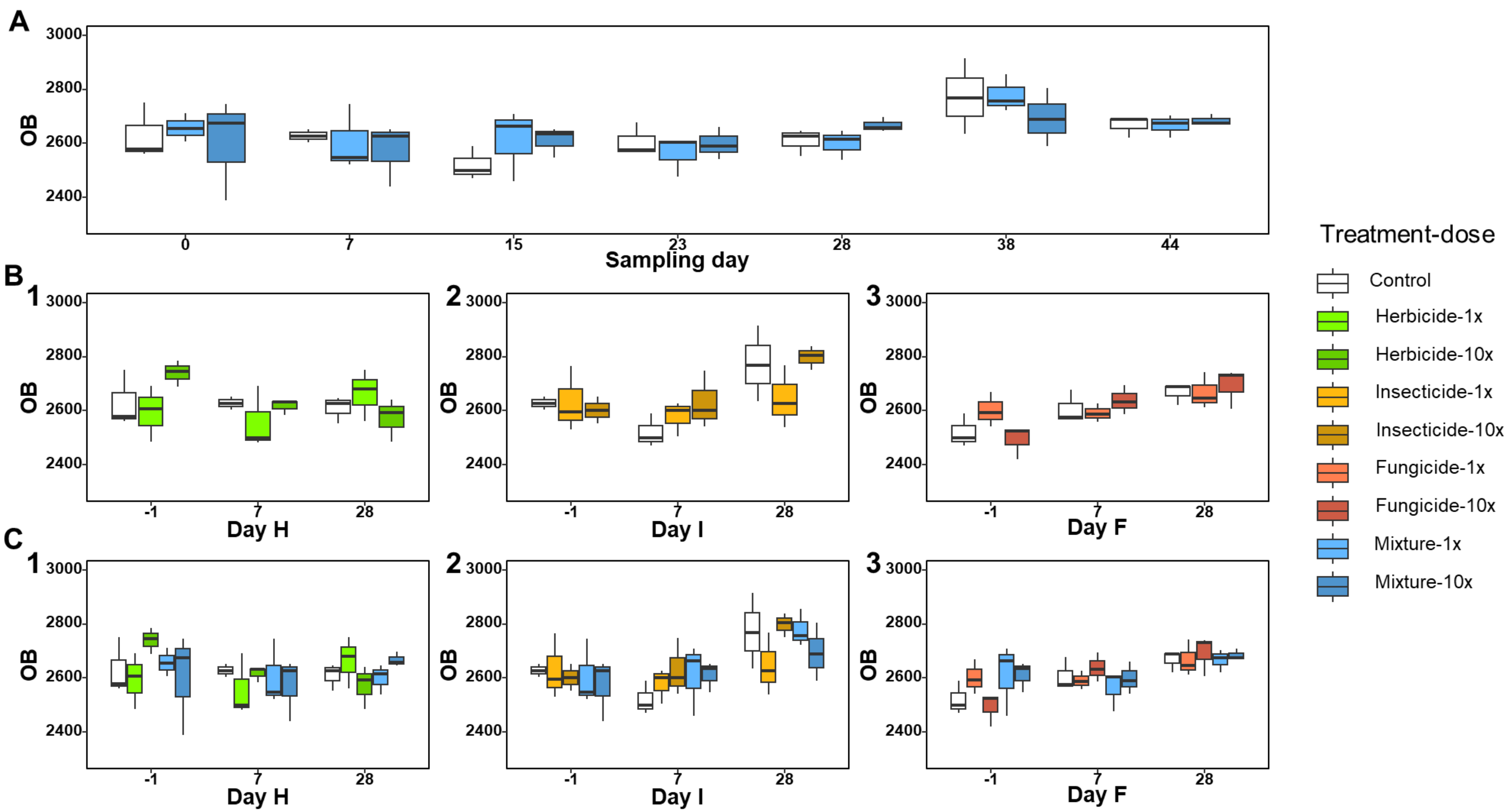
A
B
C
1
2
3
OB
3000
2800
2600
2400
0
7
15
23
28
38
44
Sampling day
-1
Day H
Day I
Day F
Treatment-dose
Control
Herbicide-1x
Herbicide-10x
Insecticide-1x
Insecticide-10x
Fungicide-1x
Fungicide-10x
Mixture-1x
Mixture-10x

Figure S5.

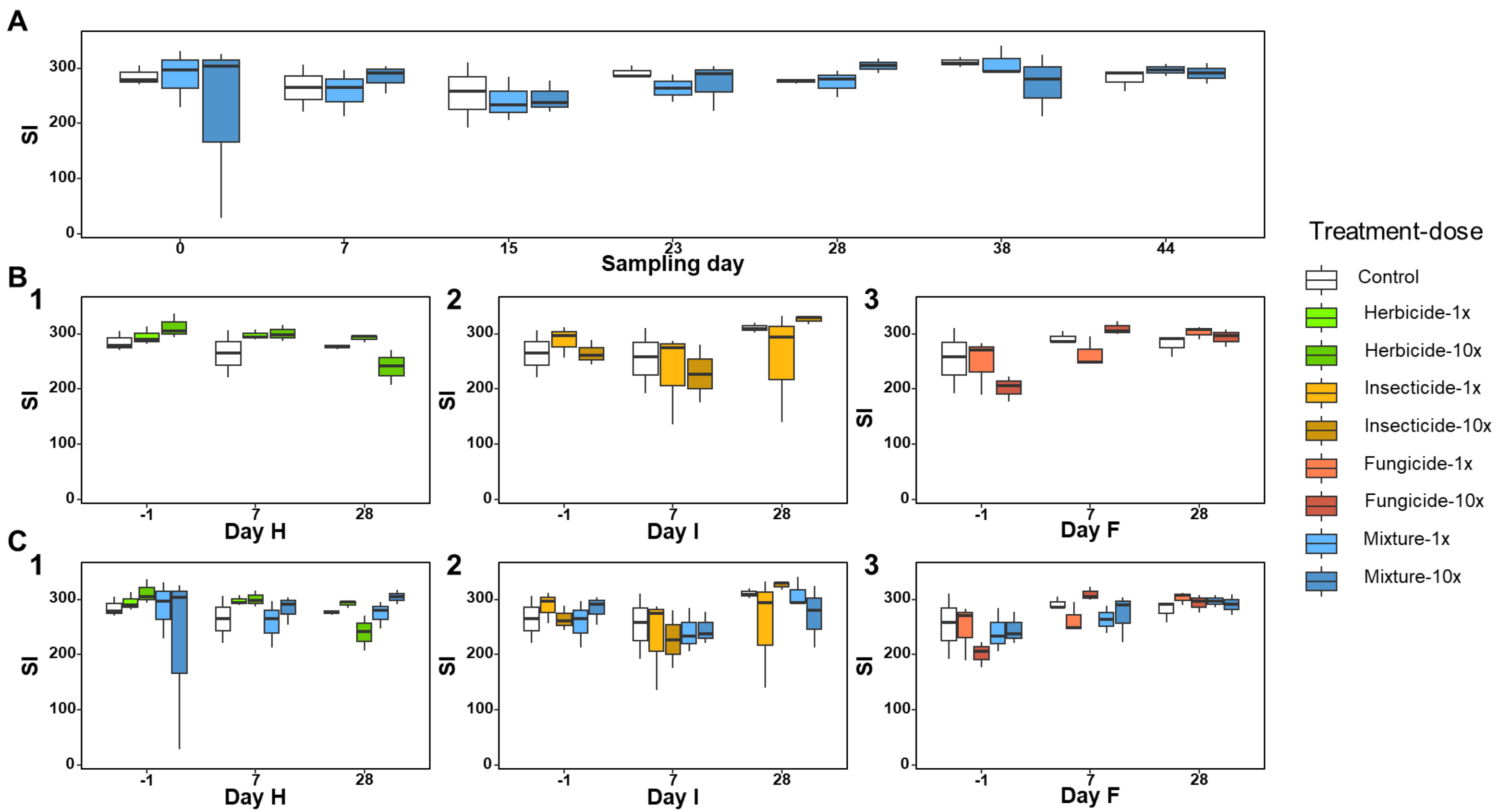

A
B
C
1
2
3
SI
Sampling day
Day H
Day I
Day F
Treatment-dose
Control
Herbicide-1x
Herbicide-10x
Insecticide-1x
Insecticide-10x
Fungicide-1x
Fungicide-10x
Mixture-1x
Mixture-10x


Figure S6.

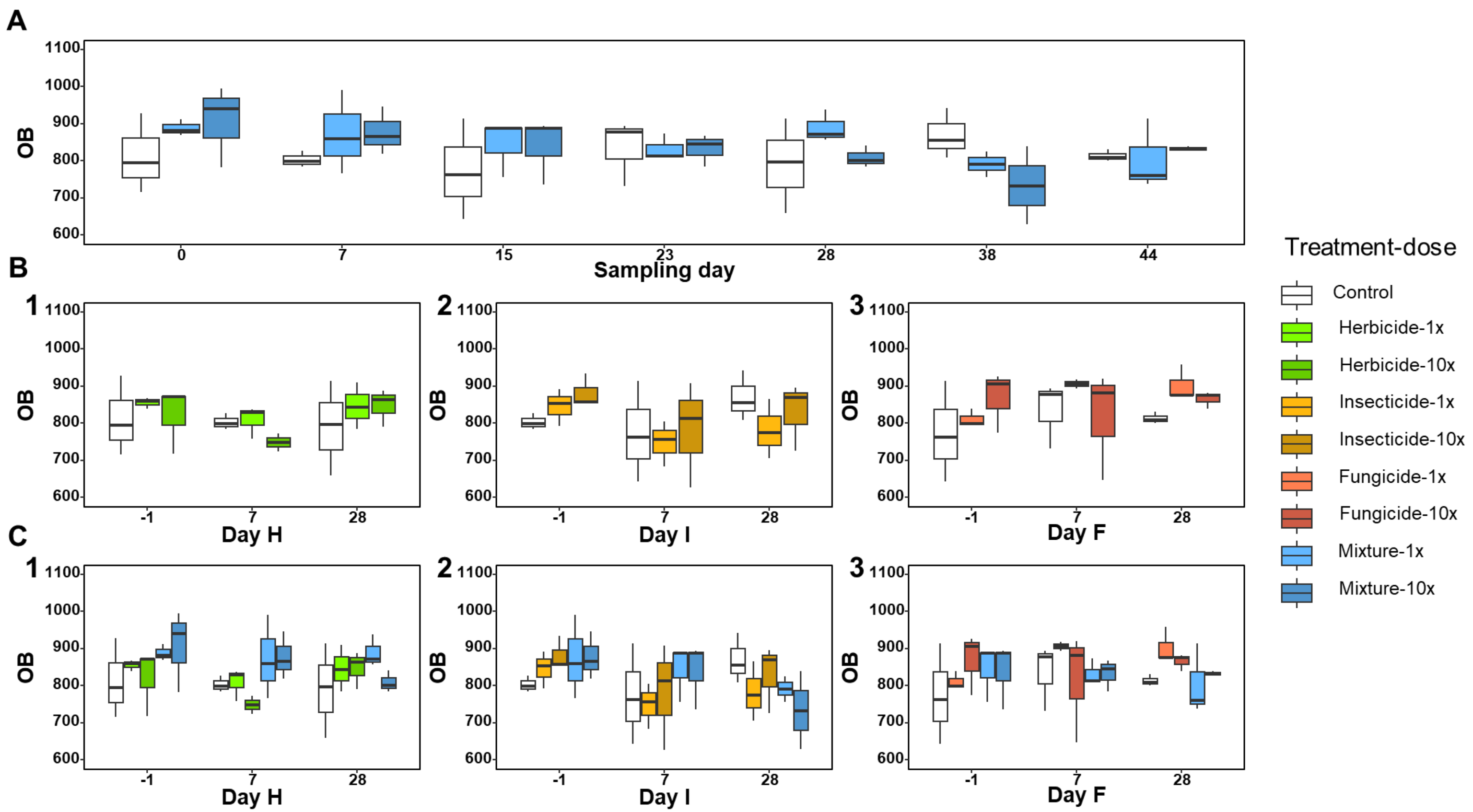
A
B
C
OB
Sampling day
Day H
Day I
Day F
Treatment-dose
Control
Herbicide-1x
Herbicide-10x
Insecticide-1x
Insecticide-10x
Fungicide-1x
Fungicide-10x
Mixture-1x
Mixture-10x

Figure S7.

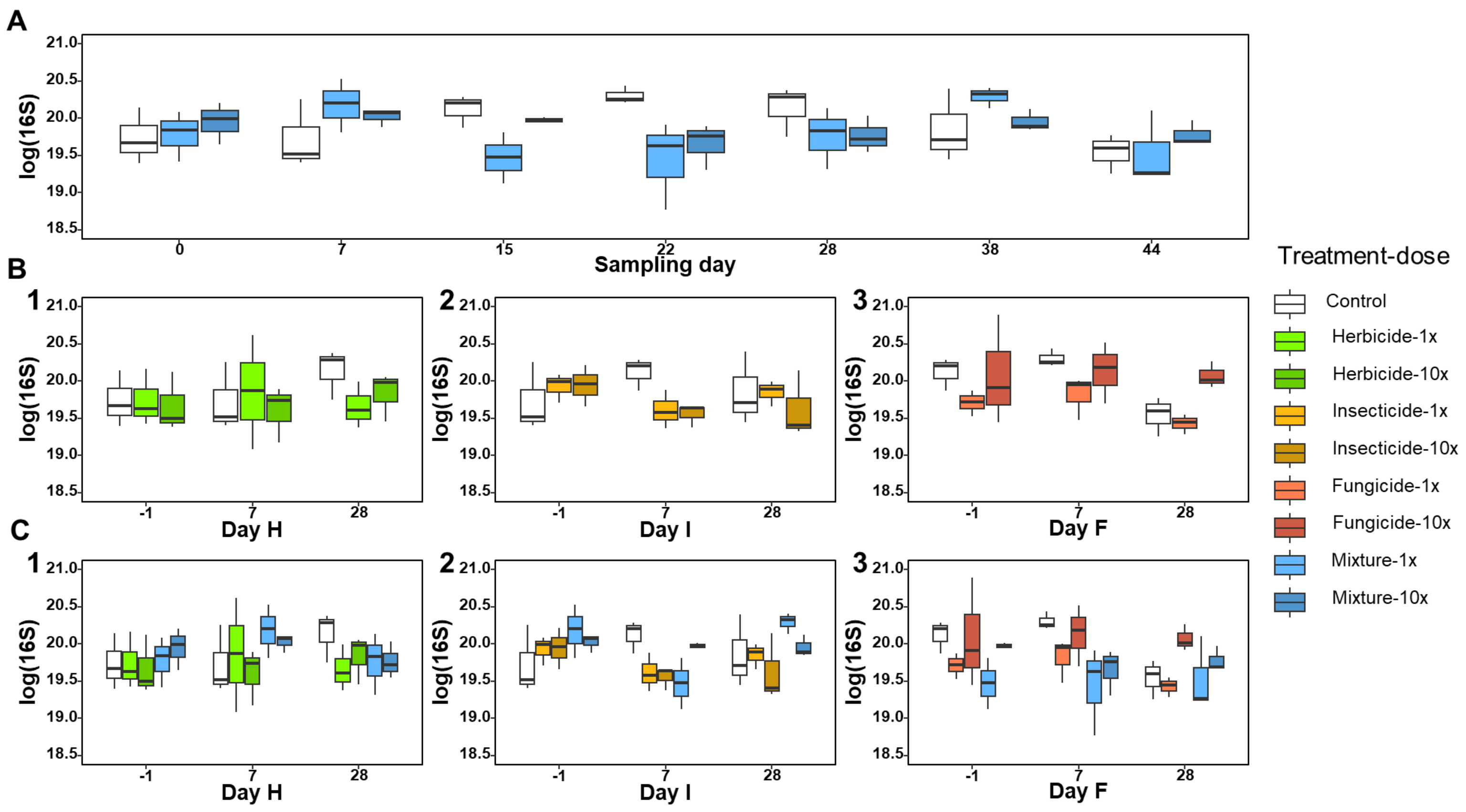
A
log(16S)
Sampling day
B
C
1
2
3
Day H
Day I
Day F
Treatment-dose
Control
Herbicide-1x
Herbicide-10x
Insecticide-1x
Insecticide-10x
Fungicide-1x
Fungicide-10x
Mixture-1x
Mixture-10x

Figure S8.

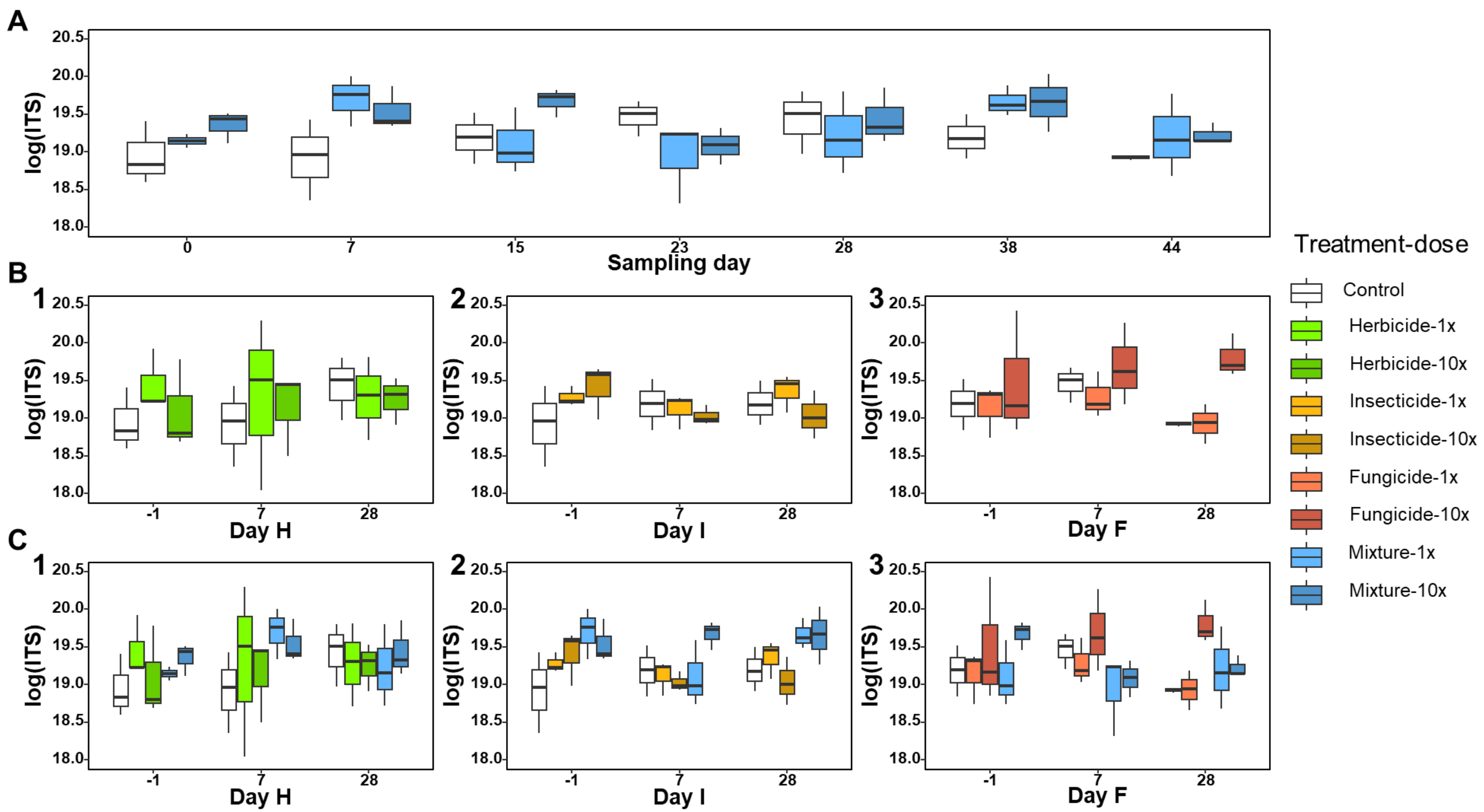

A
B
C
1
2
3
log(ITS)
20.5
20.0
19.5
19.0
18.5
18.0
0
7
15
23
28
38
44
Sampling day
-1
Day H
Day I
Day F
Treatment-dose
Control
Herbicide-1x
Herbicide-10x
Insecticide-1x
Insecticide-10x
Fungicide-1x
Fungicide-10x
Mixture-1x
Mixture-10x


Figure S9.

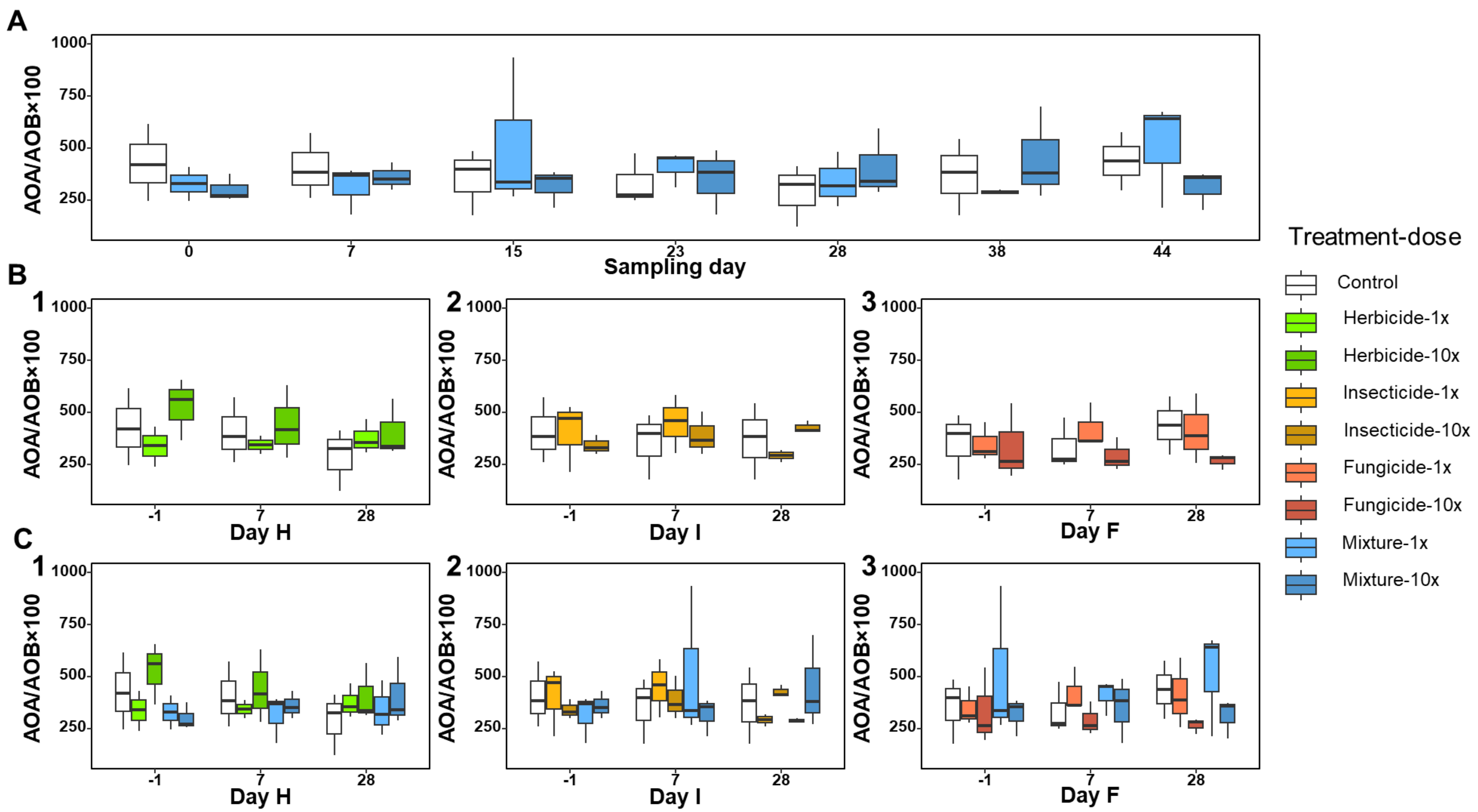
A
B
C
1
2
3
AOA/AOB×100
1000
750
500
250
0
7
15
23
28
38
44
Sampling day
-1
Day H
Day I
Day F
Treatment-dose
Control
Herbicide-1x
Herbicide-10x
Insecticide-1x
Insecticide-10x
Fungicide-1x
Fungicide-10x
Mixture-1x
Mixture-10x

Figure S10.

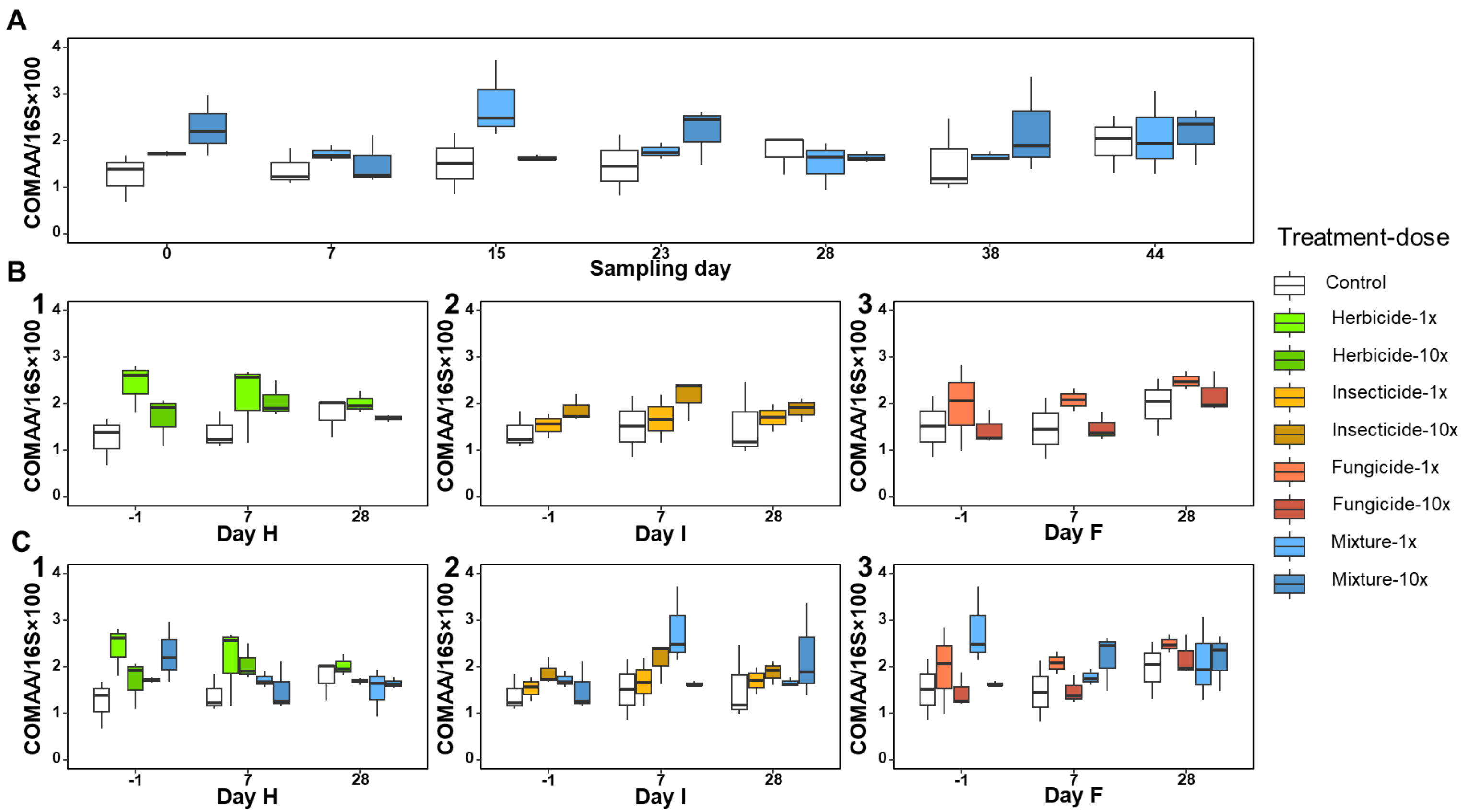

A
COMAA/16S×100
Sampling day
0
7
15
23
28
38
44
B
1
2
3
Day H
Day I
Day F
-1
7
28
C
Treatment-dose
Control
Herbicide-1x
Herbicide-10x
Insecticide-1x
Insecticide-10x
Fungicide-1x
Fungicide-10x
Mixture-1x
Mixture-10x


Figure S11.

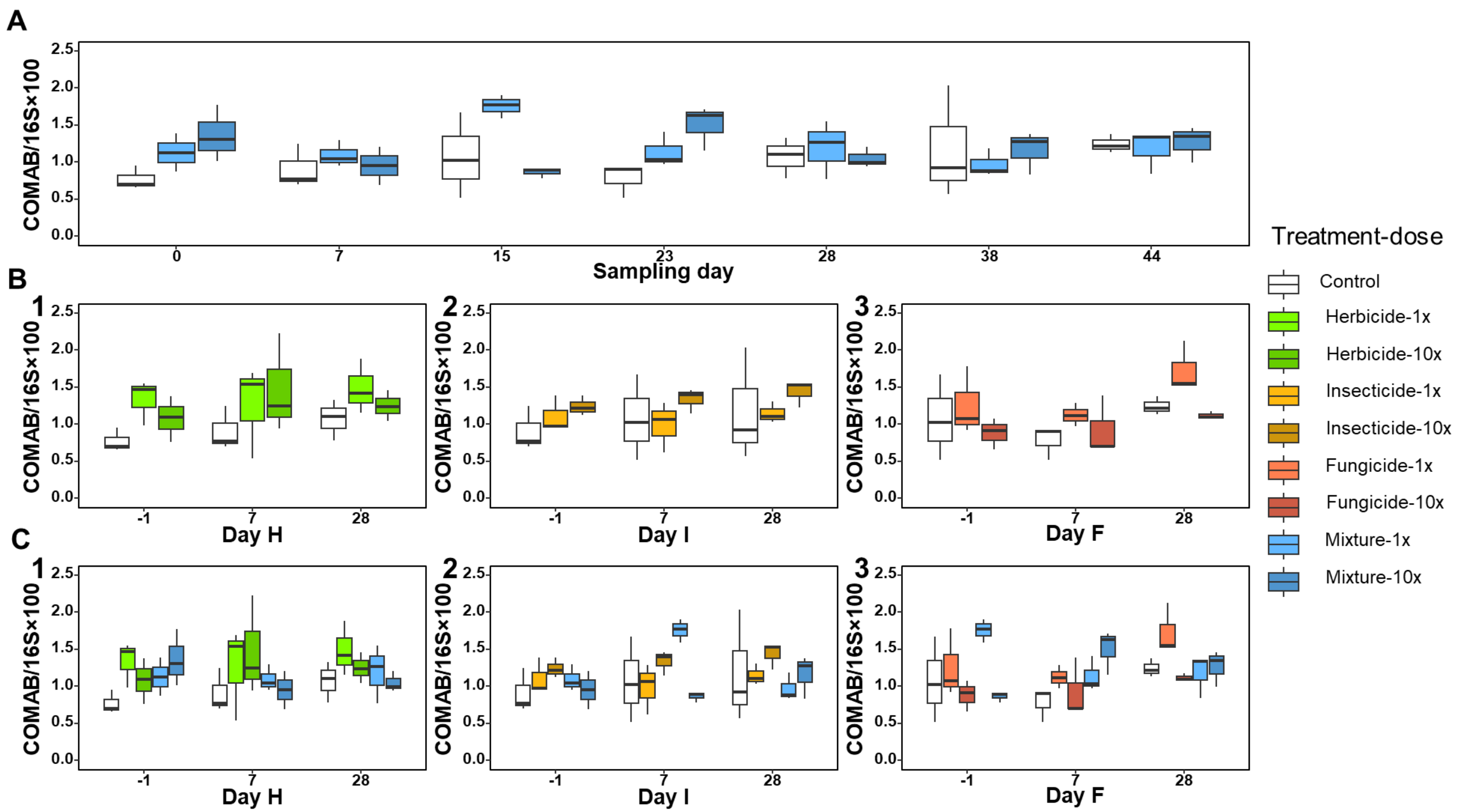
A
B
C
1
2
3
COMAB/16S×100
Sampling day
Day H
Day I
Day F
Treatment-dose
Control
Herbicide-1x
Herbicide-10x
Insecticide-1x
Insecticide-10x
Fungicide-1x
Fungicide-10x
Mixture-1x
Mixture-10x

Figure S12.

Figure S1. Schematic representation of the field plot design, with the corresponding X and Y plot coordinates

Figure S2. Pearson correlation analysis between the plot coordinates and the soil properties. CEC: cation exchange capacity, RATIOCN: ratio C/N, NTOT: total nitrogen, OM: organic material, OC: organic carbon

Figure S3. Changes in pH, OC (organic carbon), OM (organic material), NTOT (total nitrogen), CNRATIO (ratio C/N) and CEC (cation exchange capacity) according to the X and Y axis in the field site

Figure S4. 16S phylogenetic diversity (PD) for the mixture (panel A), the single pesticides (panel B, 1-2-3) and the mixture and single treatments together (panel C, 1-2-3) at the different sampling dates. Day H, Day I and Day F represent the sampling day relative to the individual pesticide application (-1: 1 day before respective application; 7: 7 days after respective application; 28: 28 days after respective application). The treatments are control, herbicide (clopyralid), insecticide (zeta-cypermethrin), fungicide (pyraclostrobin) and sequential mixture (mixture), all tested at 1× or 10× the agronomical dose. Shown are the medians, the lower and upper hinges which correspond to the first and third quartiles

Figure S5. 16S observed species (OB) for the mixture (panel A), the single pesticides (panel B, 1-2-3) and the mixture and single treatments together (panel C, 1-2-3) at the different sampling dates. Day H, Day I and Day F represent the sampling day relative to the individual pesticide application (-1: 1 day before respective application; 7: 7 days after respective application; 28: 28 days after respective application). The treatments are control, herbicide (clopyralid), insecticide (zeta-cypermethrin), fungicide (pyraclostrobin) and sequential mixture (mixture), all tested at 1× or 10× the agronomical dose. Shown are the medians, the lower and upper hinges which correspond to the first and third quartiles

Figure S6. 16S Simpson index (SI) for the mixture (panel A), the single pesticides (panel B, 1-2-3) and the mixture and single treatments together (panel C, 1-2-3) at the different sampling dates. Day H, Day I and Day F represent the sampling day relative to the individual pesticide application (-1: 1 day before respective application; 7: 7 days after respective application; 28: 28 days after respective application). The treatments are control, herbicide (clopyralid), insecticide (zeta-cypermethrin), fungicide (pyraclostrobin) and sequential mixture (mixture), all tested at 1× or 10× the agronomical dose. Shown are the medians, the lower and upper hinges which correspond to the first and third quartiles

Figure S7. ITS Observed species (OB) for the mixture (panel A), the single pesticides (panel B, 1-2-3) and the mixture and single treatments together (panel C, 1-2-3) at different sampling dates. Day H, Day I and Day F represent the sampling day relative to the individual pesticide application (-1: 1 day before respective application; 7: 7 days after respective application; 28: 28 days after respective application). The treatments are control, herbicide (clopyralid), insecticide (zeta-cypermethrin), fungicide (pyraclostrobin) and sequential mixture (mixture), all tested at 1× or 10× the agronomical dose. Shown are the medians, the lower and upper hinges which correspond to the first and third quartiles

Figure S8. 16S abundance (copy number/g of dry soil) for the mixture (panel A), the single pesticides (panel B, 1-2-3) and the mixture and single treatments together (panel C, 1-2-3) at the different sampling dates. Day H, Day I and Day F represent the sampling day relative to the individual pesticide application (-1: 1 day before respective application; 7: 7 days after respective application; 28: 28 days after respective application). The treatments are control, herbicide (clopyralid), insecticide (zeta-cypermethrin), fungicide (pyraclostrobin) and sequential mixture (mixture), all tested at 1× or 10× the agronomical dose. Shown are the medians, the lower and upper hinges which correspond to the first and third quartiles

Figure S9. ITS abundance (copy number/g of dry soil) for the mixture (panel A), the single pesticides (panel B, 1-2-3) and the mixture and single treatments together (panel C, 1-2-3) at the different sampling dates. Day H, Day I and Day F represent the sampling day relative to the individual pesticide application (-1: 1 day before respective application; 7: 7 days after respective application; 28: 28 days after respective application). The treatments are control, herbicide (clopyralid), insecticide (zeta-cypermethrin), fungicide (pyraclostrobin) and sequential mixture (mixture), all tested at 1× or 10× the agronomical dose. Shown are the medians, the lower and upper hinges which correspond to the first and third quartiles

Figure S10. Relative abundance of ammonia oxidizing archaea (AOA/AOB×100) for the mixture (panel A), the single pesticides (panel B, 1-2-3) and the mixture and single treatments together (panel C, 1-2-3) at the different sampling dates. Day H, Day I and Day F represent the sampling day relative to the individual pesticide application (-1: 1 day before respective application; 7: 7 days after respective application; 28: 28 days after respective application). The treatments are control, herbicide (clopyralid), insecticide (zeta-cypermethrin), fungicide (pyraclostrobin) and sequential mixture (mixture), all tested at 1× or 10× the agronomical dose. Shown are the medians, the lower and upper hinges which correspond to the first and third quartiles

Figure S11. Relative abundance of comammox-A (COMAA/16S×100) for the mixture (panel A), the single pesticides (panel B, 1-2-3) and the mixture and single treatments together (panel C, 1-2-3) at the different sampling dates. Day H, Day I and Day F represent the sampling day relative to the individual pesticide application (-1: 1 day before respective application; 7: 7 days after respective application; 28: 28 days after respective application). The treatments are control, herbicide (clopyralid), insecticide (zeta-cypermethrin), fungicide (pyraclostrobin) and sequential mixture (mixture), all tested at 1× or 10× the agronomical dose. Shown are the medians, the lower and upper hinges which correspond to the first and third quartiles

Figure S12. Relative abundance of comammox-B (COMAB/16S×100) for the mixture (panel A), the single pesticides (panel B, 1-2-3) and the mixture and single treatments together (panel C, 1-2-3) at the different sampling dates. Day H, Day I and Day F represent the sampling day relative to the individual pesticide application (-1: 1 day before respective application; 7: 7 days after respective application; 28: 28 days after respective application). The treatments are control, herbicide (clopyralid), insecticide (zeta-cypermethrin), fungicide (pyraclostrobin) and sequential mixture (mixture), all tested at 1× or 10× the agronomical dose. Shown are the medians, the lower and upper hinges which correspond to the first and third quartiles

Table S1.

| 1 | PD_16S | | | | OB_16S | | | | SI_16S | | | |
|---|---|---|---|---|---|---|---|---|---|---|---|---|
| | Estimate | SE | Z value | Pr(>\|z\|) | Estimate | SE | Z value | Pr(>\|z\|) | Estimate | SE | Z value | Pr(>\|z\|) |
| Mixture-1× | 0.12 | 3.48 | 0.03 | 0.97 | 4.72 | 27.14 | 0.17 | 0.86 | -16.68 | 14.55 | -1.15 | 0.25 |
| Mixture-10× | 0.02 | 3.86 | 0.01 | 1.00 | -24.99 | 30.11 | -0.83 | 0.41 | -6.18 | 16.14 | -0.38 | 0.70 |
| OM | 0.98 | 0.96 | 1.03 | 0.31 | 10.42 | 7.48 | 1.39 | 0.16 | 1.29 | 4.01 | 0.32 | 0.75 |
| pH | 6.85 | 20.84 | 0.33 | 0.74 | 53.39 | 162.52 | 0.33 | 0.74 | 84.76 | 87.14 | 0.97 | 0.33 |
| RATIOCN | 0.57 | 6.20 | 0.09 | 0.93 | 33.41 | 48.35 | 0.69 | 0.49 | 4.15 | 25.93 | 0.16 | 0.87 |
| CEC | 0.05 | 0.67 | 0.08 | 0.94 | 2.85 | 5.23 | 0.54 | 0.59 | 5.13 | 2.80 | 1.83 | 0.07 |

| 2 | PD_16S | | | | OS_16S | | | | SI_16S | | | |
|---|---|---|---|---|---|---|---|---|---|---|---|---|
| | Estimate | SE | Z value | Pr(>\|z\|) | Estimate | SE | Z value | Pr(>\|z\|) | Estimate | SE | Z value | Pr(>\|z\|) |
| Herbicide-1× | 2.47 | 4.61 | 0.54 | 0.59 | -1.23 | 34.26 | -0.04 | 0.97 | 13.94 | 15.82 | 0.88 | 0.38 |
| Herbicide-10× | 2.82 | 4.43 | 0.64 | 0.52 | 15.15 | 32.96 | 0.46 | 0.65 | 3.95 | 15.21 | 0.26 | 0.80 |
| Insecticide-1× | -1.44 | 4.65 | -0.31 | 0.76 | -15.15 | 34.49 | -0.44 | 0.66 | 28.07 | 15.99 | -1.76 | 0.08 |
| Insecticide-10× | 5.49 | 4.43 | 1.24 | 0.22 | 51.51 | 32.88 | 1.57 | 0.12 | 1.66 | 15.24 | 0.11 | 0.91 |
| Fungicide-1× | -1.49 | 4.66 | -0.32 | 0.75 | -5.74 | 34.72 | -0.17 | 0.87 | -15.34 | 16.01 | -0.96 | 0.34 |
| Fungicide-10× | -3.23 | 5.10 | -0.63 | 0.53 | -15.22 | 37.97 | -0.40 | 0.69 | -24.79 | 17.54 | -1.41 | 0.16 |
| OM | -1.10 | 0.71 | -1.56 | 0.12 | -3.56 | 5.22 | -0.68 | 0.50 | -1.95 | 2.43 | -0.80 | 0.42 |
| pH | -10.14 | 12.74 | -0.80 | 0.43 | -91.25 | 94.25 | -0.97 | 0.33 | -82.34 | 43.82 | -1.88 | 0.06 |
| RATIOCN | 0.74 | 5.29 | 0.14 | 0.89 | -8.04 | 39.17 | -0.21 | 0.84 | 7.97 | 18.21 | 0.44 | 0.66 |
| CEC | -0.81 | 0.52 | -1.55 | 0.12 | -4.62 | 3.88 | -1.19 | 0.23 | 1.42 | 1.80 | 0.79 | 0.43 |

<table>
<tr><td rowspan="2">3</td><td></td><td colspan="3">PD_16S</td><td colspan="4">OB_16S</td><td colspan="4">SI_16S</td></tr>
<tr><td>Estimate</td><td>SE</td><td>Z value</td><td>Pr(>|z|)</td><td>Estimate</td><td>SE</td><td>Z value</td><td>Pr(>|z|)</td><td>Estimate</td><td>SE</td><td>Z value</td><td>Pr(>|z|)</td></tr>
<tr><td>Herbicide-1×</td><td>1.62</td><td>5.04</td><td>0.32</td><td>0.75</td><td>6.31</td><td>42.31</td><td>0.15</td><td>0.88</td><td>28.98</td><td>22.06</td><td>1.31</td><td>0.19</td></tr>
<tr><td>Herbicide-10×</td><td>2.05</td><td>4.55</td><td>0.45</td><td>0.65</td><td>16.28</td><td>38.15</td><td>0.43</td><td>0.67</td><td>4.26</td><td>19.90</td><td>0.21</td><td>0.83</td></tr>
<tr><td>Mixture-1×</td><td>-4.87</td><td>4.87</td><td>-1.00</td><td>0.32</td><td>-12.83</td><td>40.87</td><td>-0.31</td><td>0.75</td><td>-10.73</td><td>21.31</td><td>-0.50</td><td>0.61</td></tr>
<tr><td>Mixture-10×</td><td>6.29</td><td>5.06</td><td>1.24</td><td>0.21</td><td>35.46</td><td>42.43</td><td>0.84</td><td>0.40</td><td>20.89</td><td>22.13</td><td>0.94</td><td>0.35</td></tr>
<tr><td>OM</td><td>-0.90</td><td>0.86</td><td>-1.05</td><td>0.29</td><td>-1.05</td><td>7.19</td><td>-0.15</td><td>0.88</td><td>-0.87</td><td>3.75</td><td>-0.23</td><td>0.82</td></tr>
<tr><td>pH</td><td>40.0</td><td>22.37</td><td>1.79</td><td>0.07</td><td>204.45</td><td>187.68</td><td>1.09</td><td>0.28</td><td>-18.94</td><td>97.87</td><td>-0.19</td><td>0.85</td></tr>
<tr><td>RATIOCN</td><td>-2.78</td><td>7.09</td><td>-0.39</td><td>0.69</td><td>-46.42</td><td>59.47</td><td>-0.78</td><td>0.44</td><td>-59.83</td><td>31.01</td><td>-1.93</td><td>0.05</td></tr>
<tr><td>CEC</td><td>-0.44</td><td>0.75</td><td>-0.59</td><td>0.55</td><td>-4.66</td><td>6.32</td><td>-0.74</td><td>0.46</td><td>3.95</td><td>3.29</td><td>1.20</td><td>0.23</td></tr>
</table>

<table>
<tr><td rowspan="2">4</td><td colspan="4">PD_16S</td><td colspan="4">OB_16S</td><td colspan="4">SI_16S</td></tr>
<tr><td>Estimate</td><td>SE</td><td>Z value</td><td>Pr(>|z|)</td><td>Estimate</td><td>SE</td><td>Z value</td><td>Pr(>|z|)</td><td>Estimate</td><td>SE</td><td>Z value</td><td>Pr(>|z|)</td></tr>
<tr><td>Insecticide-1×</td><td>-2.24</td><td>5.52</td><td>-0.41</td><td>0.68</td><td>-39.59</td><td>40.98</td><td>-0.97</td><td>0.33</td><td>-35.64</td><td>20.94</td><td>-1.70</td><td>0.09</td></tr>
<tr><td>Insecticide-10×</td><td>6.23</td><td>5.39</td><td>1.16</td><td>0.25</td><td>51.96</td><td>40.02</td><td>1.30</td><td>0.19</td><td>15.82</td><td>20.45</td><td>0.77</td><td>0.44</td></tr>
<tr><td>Mixture-1×</td><td>7.15</td><td>5.20</td><td>1.38</td><td>0.17</td><td>47.61</td><td>38.63</td><td>1.23</td><td>0.22</td><td>5.07</td><td>19.74</td><td>0.26</td><td>0.80</td></tr>
<tr><td>Mixture-10×</td><td>-1.35</td><td>5.70</td><td>-0.24</td><td>0.81</td><td>-64.36</td><td>42.37</td><td>-1.52</td><td>0.13</td><td>-32.03</td><td>21.65</td><td>-1.48</td><td>0.14</td></tr>
<tr><td>OM</td><td>-0.07</td><td>1.27</td><td>-0.06</td><td>0.95</td><td>0.50</td><td>9.45</td><td>0.05</td><td>0.96</td><td>5.47</td><td>4.83</td><td>1.13</td><td>0.26</td></tr>
<tr><td>pH</td><td>-12.74</td><td>21.70</td><td>-0.59</td><td>0.56</td><td>-139.67</td><td>161.20</td><td>-0.87</td><td>0.39</td><td>-215.70</td><td>82.38</td><td>-2.62</td><td>0.01</td></tr>
<tr><td>RATIOCN</td><td>5.46</td><td>6.81</td><td>0.80</td><td>0.42</td><td>76.95</td><td>50.61</td><td>1.52</td><td>0.13</td><td>-14.09</td><td>25.87</td><td>-0.55</td><td>0.59</td></tr>
<tr><td>CEC</td><td>-1.06</td><td>0.75</td><td>-1.41</td><td>0.16</td><td>-6.43</td><td>5.56</td><td>-1.16</td><td>0.25</td><td>2.35</td><td>2.84</td><td>0.83</td><td>0.41</td></tr>
</table>

| 5 | PD_16S | | | | OB_16S | | | | SI_16S | | | |
|---|---|---|---|---|---|---|---|---|---|---|---|---|
| | Estimate | SE | Z value | Pr(>\|z\|) | Estimate | SE | Z value | Pr(>\|z\|) | Estimate | SE | Z value | Pr(>\|z\|) |
| Fungicide-1× | 2.76 | 4.75 | 0.58 | 0.56 | 23.11 | 30.85 | 0.75 | 0.45 | -1.73 | 13.57 | -0.13 | 0.90 |
| Fungicide-10× | -2.51 | 5.55 | -0.45 | 0.65 | -10.58 | 36.09 | -0.29 | 0.77 | -1.96 | 15.87 | -0.12 | 0.90 |
| Mixture-1× | 3.86 | 4.62 | 0.84 | 0.40 | 26.03 | 30.04 | 0.87 | 0.39 | -16.78 | 13.21 | -1.27 | 0.20 |
| Mixture-10× | 3.17 | 4.94 | 0.64 | 0.52 | 18.42 | 32.09 | 0.57 | 0.57 | -1.77 | 14.11 | -0.13 | 0.90 |
| OM | 0.71 | 1.12 | 0.63 | 0.53 | 9.22 | 7.31 | 1.26 | 0.21 | -0.32 | 3.21 | -0.10 | 0.92 |
| pH | -8.34 | 13.13 | -0.64 | 0.53 | -49.65 | 85.30 | -0.58 | 0.56 | 53.70 | 37.51 | 1.43 | 0.15 |
| RATIOCN | -1.98 | 4.74 | -0.42 | 0.68 | -5.17 | 30.82 | -0.17 | 0.87 | 2.86 | 13.55 | 0.21 | 0.83 |
| CEC | -0.49 | 0.65 | -0.76 | 0.45 | -3.56 | 4.23 | -0.84 | 0.40 | 3.79 | 1.86 | 2.04 | **0.04** |

Table S2.

| 1 | OS_ITS | | | | SI_ITS | | | |
|---|---|---|---|---|---|---|---|---|
| | Estimate | SE | Z value | Pr(>\|z\|) | Estimate | SE | Z value | Pr(>\|z\|) |
| ixture-1× | 47.28 | 24.40 | 1.94 | 0.05 | 6.97 | 2.71 | 2.58 | **0.01** |
| Mixture-10× | -7.20 | 26.95 | -0.27 | 0.79 | 1.94 | 2.99 | 0.65 | 0.52 |
| OM | 21.72 | 6.77 | 3.21 | **<0.01** | 1.85 | 0.75 | 2.47 | **0.01** |
| pH | -137.16 | 145.25 | -0.94 | 0.34 | -6.86 | 16.10 | -0.43 | 0.67 |
| RATIOCN | -37.12 | 44.22 | -0.84 | 0.40 | 0.68 | 4.90 | 0.14 | 0.89 |
| CEC | 2.58 | 4.76 | 0.54 | 0.59 | -0.90 | 0.53 | -1.70 | 0.09 |

| Pairwise comparison-1 | SI_ITS | | | |
|---|---|---|---|---|
| | Emmean | SE | df | group |
| Control | 20.70 | 1.80 | 52 | B |
| Mixture-1× | 27.70 | 2.05 | 52 | A |
| Mixture-10× | 22.70 | 2.18 | 52 | AB |

| 2 | OB_ITS | | | | SI_ITS | | | |
|---|---|---|---|---|---|---|---|---|
| | Estimate | SE | Z value | Pr(>\|z\|) | Estimate | SE | Z value | Pr(>\|z\|) |
| Herbicide-1× | 14.11 | 29.34 | 0.48 | 0.63 | 8.24 | 3.18 | 2.59 | **0.01** |
| Herbicide-10× | 7.86 | 29.27 | 0.27 | 0.79 | 9.97 | 3.17 | 3.15 | **<0.01** |
| Insecticide-1× | -13.69 | 30.13 | -0.45 | 0.65 | 4.74 | 3.26 | 1.45 | 0.15 |
| Insecticide-10× | 28.71 | 28.56 | 1.01 | 0.31 | 4.08 | 3.09 | 1.32 | 0.19 |
| Fungicide-1× | 62.62 | 30.72 | 2.04 | **0.04** | 6.32 | 3.33 | 1.90 | 0.06 |
| Fungicide-10× | 20.51 | 33.05 | 0.62 | 0.53 | 2.49 | 3.58 | 0.70 | 0.49 |
| OM | 10.09 | 4.80 | 2.10 | **0.04** | 0.49 | 0.52 | 0.95 | 0.34 |
| pH | -20.45 | 87.16 | -0.24 | 0.81 | -10.34 | 9.44 | -1.10 | 0.27 |
| RATIOCN | 2.66 | 37.08 | 0.07 | 0.94 | 1.13 | 4.02 | 0.28 | 0.78 |
| CEC | -5.24 | 3.62 | -1.45 | 0.15 | -1.32 | 0.39 | -3.36 | **<0.01** |

| Pairwise comparison-2 | OB_ITS | | | | SI_ITS | | | |
|---|---|---|---|---|---|---|---|---|
| | Emmean | SE | df | group | Emmean | SE | df | group |
| Control | 813 | 15.80 | 60 | A | 20.1 | 1.71 | 60 | B |
| Herbicide-1× | 827 | 24.90 | 60 | A | 28.4 | 2.70 | 60 | AB |
| Herbicide-10× | 821 | 25.30 | 60 | A | 30.1 | 2.74 | 60 | A |
| Insecticide-1× | 799 | 24.50 | 60 | A | 24.9 | 2.66 | 60 | AB |
| Insecticide-10× | 841 | 25.20 | 60 | A | 24.2 | 2.73 | 60 | AB |
| Fungicide-1× | 875 | 25.50 | 60 | A | 26.5 | 2.76 | 60 | AB |
| Fungicide-10× | 833 | 27.30 | 60 | A | 22.6 | 2.96 | 60 | AB |

| 3 | OB_ITS | | | | SI_ITS | | | |
|---|---|---|---|---|---|---|---|---|
| | Estimate | SE | Z value | Pr(>\|z\|) | Estimate | SE | Z value | Pr(>\|z\|) |
| Herbicide-1× | 53.45 | 31.91 | 1.68 | 0.09 | 10.24 | 3.31 | 3.10 | <0.01 |
| Herbicide-10× | 12.32 | 29.62 | 0.42 | 0.68 | 8.42 | 3.07 | 2.74 | 0.01 |
| Mixture-1× | 88.32 | 30.87 | 2.86 | <0.01 | 7.82 | 3.20 | 2.44 | 0.01 |
| Mixture-10× | 89.36 | 31.94 | 2.80 | 0.01 | 11.22 | 3.31 | 3.39 | <0.01 |
| OM | 8.69 | 5.45 | 1.60 | 0.11 | 0.68 | 0.56 | 1.20 | 0.23 |
| pH | -112.38 | 145.80 | -0.77 | 0.44 | -22.22 | 15.11 | -1.47 | 0.14 |
| RATIOCN | -108.07 | 45.57 | -2.37 | 0.02 | -10.48 | 4.72 | -2.22 | 0.03 |
| CEC | -2.99 | 4.78 | -0.63 | 0.53 | -1.48 | 0.50 | -2.99 | <0.01 |

| Pairwise comparison-3 | OB_ITs | | | | SI_ITS | | | |
|---|---|---|---|---|---|---|---|---|
| | Emmean | SE | df | group | Emmean | SE | df | group |
| Control | 791 | 21.00 | 33 | A | 19.6 | 2.18 | 33 | B |
| Herbicide-1× | 844 | 22.00 | 33 | A | 29.8 | 2.28 | 33 | A |
| Herbicide-10× | 803 | 22.60 | 33 | A | 28.0 | 2.34 | 33 | AB |
| Mixture-1× | 879 | 22.20 | 33 | A | 27.4 | 2.30 | 33 | AB |
| Mixture-10× | 880 | 23.50 | 33 | A | 30.8 | 2.43 | 33 | A |

| 4 | OB_ITS | | | | SI_ITS | | | |
|---|---|---|---|---|---|---|---|---|
| | Estimate | SE | Z value | Pr(>\|z\|) | Estimate | SE | Z value | Pr(>\|z\|) |
| Insecticide-1× | -22.39 | 39.92 | -0.56 | 0.57 | 4.63 | 4.12 | 1.13 | 0.26 |
| Insecticide-10× | 44.60 | 39.13 | 1.14 | 0.25 | 3.08 | 4.04 | 0.76 | 0.44 |
| Mixture-1× | 44.86 | 38.83 | 1.16 | 0.25 | 7.98 | 4.00 | 1.99 | **0.05** |
| Mixture-10× | -18.27 | 41.29 | -0.44 | 0.66 | -1.10 | 4.26 | -0.26 | 0.80 |
| OM | 20.01 | 9.50 | 2.11 | **0.04** | 0.60 | 0.98 | 0.61 | 0.54 |
| pH | -183.87 | 158.27 | -1.16 | 0.25 | -3.59 | 16.32 | -0.22 | 0.83 |
| RATIOCN | -56.76 | 49.99 | -1.14 | 0.26 | 0.63 | 5.15 | 0.12 | 0.90 |
| CEC | 0.03 | 5.56 | 0.01 | 1.00 | -1.02 | 0.57 | -1.78 | 0.07 |

| Pairwise comparison-4 | SI_ITS | | | |
|---|---|---|---|---|
| | Emmeans | SE | df | group |
| Control | 20.9 | 2.70 | 33 | A |
| Insecticide-1× | 25.5 | 3.01 | 33 | A |
| Insecticide-10× | 23.9 | 3.07 | 33 | A |
| Mixture-1× | 28.8 | 3.00 | 33 | A |
| Mixture-10× | 19.8 | 3.20 | 33 | A |

| 5 | OB_ITS | | | | SI_ITS | | | |
|---|---|---|---|---|---|---|---|---|
| | Estimate | SE | Z value | Pr(>\|z\|) | Estimate | SE | Z value | Pr(>\|z\|) |
| Fungicide-1× | 72.32 | 29.74 | 2.43 | **0.02** | 10.28 | 4.07 | 2.53 | **0.01** |
| Fungicide-10× | 0.96 | 33.96 | 0.03 | 0.98 | 2.52 | 4.64 | 0.54 | 0.59 |
| Mixture-1× | 35.38 | 28.27 | 1.25 | 0.21 | 10.75 | 3.87 | 2.78 | **0.01** |
| Mixture-10× | -2.14 | 30.39 | -0.07 | 0.94 | 2.86 | 4.16 | 0.69 | 0.49 |
| OM | 23.13 | 6.86 | 3.37 | **<0.01** | 2.29 | 0.94 | 2.44 | **0.01** |
| pH | -33.54 | 80.29 | -0.42 | 0.68 | -11.47 | 10.98 | -1.05 | 0.30 |
| RATIOCN | -0.71 | 30.09 | -0.02 | 0.98 | 0.85 | 4.11 | 0.21 | 0.84 |
| CEC | -4.52 | 4.02 | -1.12 | 0.26 | -1.41 | 0.55 | -2.56 | **0.01** |

| Pairwise comparison-5 | OB_ITS | | | | SI_ITS | | | |
|---|---|---|---|---|---|---|---|---|
| | Emmean | SE | df | group | Emmean | SE | df | group |
| Control | 816 | 20.50 | 32 | A | 17.8 | 2.80 | 32 | A |
| Fungicide-1× | 889 | 21.30 | 32 | A | 28.1 | 2.91 | 32 | A |
| Fungicide-10× | 817 | 24.00 | 32 | A | 20.3 | 3.28 | 32 | A |
| Mixture-1× | 852 | 22.30 | 32 | A | 28.6 | 3.04 | 32 | A |
| Mixture-10× | 814 | 21.60 | 32 | A | 20.7 | 2.96 | 32 | A |

Table S3.

| 1a | 16S+ | | | | ITS+ | | | | AOA/AOB×100 | | | |
|---|---|---|---|---|---|---|---|---|---|---|---|---|
| | Estimate | SE | Z value | Pr(>\|z\|) | Estimate | SE | Z value | Pr(>\|z\|) | Estimate | SE | Z value | Pr(>\|z\|) |
| Mixture-1× | -0.15 | 0.13 | -1.21 | 0.22 | 0.10 | 0.13 | 0.82 | 0.41 | 86.67 | 41.36 | 2.10 | **0.04** |
| Mixture-10× | -0.01 | 0.14 | -0.09 | 0.93 | 0.34 | 0.14 | 2.41 | **0.02** | -79.44 | 45.68 | -1.74 | 0.08 |
| OM | 0.01 | 0.03 | 0.03 | 0.98 | -0.01 | 0.04 | -0.22 | 0.83 | 17.70 | 11.47 | 1.54 | 0.12 |
| pH | -0.02 | 0.75 | -0.03 | 0.98 | -0.03 | 0.75 | -0.04 | 0.97 | -472.00 | 245.99 | -1.92 | 0.06 |
| RATIOCN | -0.08 | 0.22 | -0.34 | 0.73 | -0.12 | 0.22 | -0.52 | 0.60 | -18.90 | 73.17 | -0.26 | 0.80 |
| CEC | 0.01 | 0.02 | 0.34 | 0.73 | 0.02 | 0.02 | 0.92 | 0.36 | -29.57 | 8.07 | -3.66 | **<0.01** |

| 1b | AOB/16S×100 | | | | COMAA/16S×100 | | | | COMAB/16S×100 | | | |
|---|---|---|---|---|---|---|---|---|---|---|---|---|
| | Estimate | SE | Z value | Pr(>\|z\|) | Estimate | SE | Z value | Pr(>\|z\|) | Estimate | SE | Z value | Pr(>\|z\|) |
| Mixture-1× | -0.23 | 0.10 | -2.40 | **0.02** | 0.44 | 0.19 | 2.32 | **0.02** | 0.29 | 0.11 | 2.54 | **0.01** |
| Mixture-10× | 0.20 | 0.11 | 1.88 | 0.06 | 0.31 | 0.21 | 1.45 | 0.15 | 0.09 | 0.12 | 0.69 | 0.49 |
| OM | -0.06 | 0.03 | -2.23 | **0.03** | 0.04 | 0.05 | 0.75 | 0.45 | 0.05 | 0.03 | 1.69 | 0.09 |
| pH | 1.92 | 0.58 | 3.33 | **<0.01** | 2.05 | 1.14 | -1.79 | 0.07 | -0.98 | 0.67 | -1.46 | 0.14 |
| RATIOCN | 0.30 | 0.17 | 1.77 | 0.08 | -0.37 | 0.34 | -1.10 | 0.27 | -0.21 | 0.20 | -1.05 | 0.30 |
| CEC | 0.09 | 0.02 | 4.55 | **<0.01** | 0.02 | 0.04 | 0.63 | 0.53 | -0.01 | 0.02 | -0.20 | 0.84 |

| Pairwise comparison-1a | ITS | | | | AOA/AOB×100 | | | | AOB/16S×100 | | | |
|---|---|---|---|---|---|---|---|---|---|---|---|---|
| | Emmean | SE | df | group | Emmean | SE | df | group | Emmean | SE | df | group |
| Control | 19.1 | 0.08 | 53 | A | 371 | 27.70 | 53 | AB | 0.86 | 0.07 | 53 | AB |
| Mixture-1× | 19.2 | 0.10 | 53 | A | 458 | 31.60 | 53 | A | 0.63 | 0.07 | 53 | B |
| Mixture-10× | 19.5 | 0.10 | 53 | A | 292 | 32.90 | 53 | B | 1.06 | 0.08 | 53 | A |

| Pairwise comparison-1b | COMAA/16S×100 | | | | COMAB/16S×100 | | | |
|---|---|---|---|---|---|---|---|---|
| | Emmean | SE | df | group | Emmean | SE | df | group |
| Control | 1.55 | 0.13 | 53 | A | 1.00 | 0.08 | 53 | A |
| Mixture-1× | 2.00 | 0.15 | 53 | A | 1.29 | 0.09 | 53 | B |
| Mixture-10× | 1.86 | 0.15 | 53 | A | 1.09 | 0.09 | 53 | AB |

| 2a | 16S+ | | | | ITS+ | | | | AOA/AOB×100 | | | |
|---|---|---|---|---|---|---|---|---|---|---|---|---|
| | Estimate | SE | Z value | Pr(>\|z\|) | Estimate | SE | Z value | Pr(>\|z\|) | Estimate | SE | Z value | Pr(>\|z\|) |
| Herbicide-1× | -0.28 | 0.13 | -2.07 | **0.04** | 0.03 | 0.15 | 0.20 | 0.84 | -0.07 | 43.96 | -0.07 | 0.95 |
| Herbicide-10× | -0.19 | 0.13 | -1.50 | 0.13 | 0.04 | 0.14 | 0.29 | 0.78 | 2.25 | 42.03 | 2.25 | **0.02** |
| Insecticide-1× | -0.10 | 0.14 | -0.77 | 0.44 | 0.05 | 0.15 | 0.31 | 0.75 | 0.63 | 45.11 | 0.63 | 0.53 |
| Insecticide-10× | -0.15 | 0.13 | -1.13 | 0.26 | 0.11 | 0.15 | 0.73 | 0.47 | 0.45 | 42.78 | 0.45 | 0.65 |
| Fungicide-1× | -0.27 | 0.13 | -1.98 | **0.05** | -0.10 | 0.15 | -0.68 | 0.50 | 0.85 | 44.59 | 0.85 | 0.40 |
| Fungicide-10× | 0.05 | 0.15 | 0.34 | 0.73 | 0.25 | 0.17 | 1.50 | 0.13 | -1.49 | 49.34 | -1.49 | 0.14 |
| OM | 0.05 | 0.02 | 2.48 | **0.01** | 0.05 | 0.02 | 1.96 | 0.05 | 1.71 | 7.17 | 1.71 | 0.09 |
| pH | -0.17 | 0.39 | -0.45 | 0.66 | -0.63 | 0.44 | -1.43 | 0.15 | 0.02 | 129.50 | 0.02 | 0.99 |
| RATIOCN | -0.13 | 0.16 | -0.82 | 0.41 | -0.05 | 0.18 | -0.25 | 0.80 | 0.14 | 53.81 | 0.14 | 0.89 |
| CEC | 0.02 | 0.02 | 1.42 | 0.15 | 0.04 | 0.02 | 2.19 | **0.03** | -3.69 | 5.33 | -3.69 | **<0.01** |

| 2b | AOB/16S×100 | | | | COMAA/16S×100 | | | | COMAB/16S×100 | | | |
|---|---|---|---|---|---|---|---|---|---|---|---|---|
| | Estimate | SE | Z value | Pr(>\|z\|) | Estimate | SE | Z value | Pr(>\|z\|) | Estimate | SE | Z value | Pr(>\|z\|) |
| Herbicide-1× | -0.09 | 0.13 | -0.70 | 0.49 | 0.69 | 0.19 | 3.62 | **<0.00** | 0.42 | 0.14 | 3.08 | **<0.01** |
| Herbicide-10× | -0.11 | 0.12 | -0.93 | 0.35 | 0.24 | 0.18 | 1.32 | 0.19 | 0.27 | 0.13 | 2.04 | **0.04** |
| Insecticide-1× | -0.13 | 0.13 | -1.04 | 0.30 | 0.07 | 0.20 | 0.37 | 0.71 | 0.09 | 0.14 | 0.65 | 0.52 |
| Insecticide-10× | -0.16 | 0.12 | -1.31 | 0.19 | 0.37 | 0.19 | 1.99 | **0.05** | 0.29 | 0.13 | 2.21 | **0.03** |
| Fungicide-1× | -0.01 | 0.13 | -0.06 | 0.95 | 0.62 | 0.20 | 3.17 | **<0.01** | 0.39 | 0.14 | 2.82 | **<0.01** |
| Fungicide-10× | 0.25 | 0.14 | 1.80 | 0.07 | 0.24 | 0.22 | 1.10 | 0.27 | 0.06 | 0.15 | 0.38 | 0.70 |
| OM | -0.05 | 0.02 | -2.35 | **0.02** | -0.04 | 0.03 | -1.21 | 0.23 | -0.02 | 0.02 | -0.80 | 0.43 |
| pH | 0.23 | 0.37 | 0.64 | 0.53 | 0.17 | 0.55 | 0.31 | 0.76 | 0.23 | 0.39 | 0.60 | 0.55 |
| RATIOCN | -0.08 | 0.15 | -0.55 | 0.59 | 0.03 | 0.23 | 0.13 | 0.89 | 0.01 | 0.16 | 0.06 | 0.95 |
| CEC | 0.05 | 0.02 | 3.19 | **<0.01** | 0.01 | 0.02 | 0.11 | 0.91 | -0.01 | 0.02 | -0.68 | 0.50 |

| Pairwise comparison-2 | 16S | | | | AOA/AOB×100 | | | | COMAA/16S×100 | | | | COMAB/16S | | | |
|---|---|---|---|---|---|---|---|---|---|---|---|---|---|---|---|---|
| | Emmean | SE | df | group | Emmean | SE | df | group | Emmean | SE | df | group | Emmean | SE | df | group |
| Control | 19.9 | 0.07 | 62 | A | 364 | 23.80 | 62 | A | 1.55 | 0.10 | 62 | B | 0.99 | 0.08 | 62 | B |
| Herbicide-1× | 19.7 | 0.11 | 62 | A | 361 | 37.40 | 62 | A | 2.24 | 0.17 | 62 | A | 1.41 | 0.12 | 62 | A |
| Herbicide-10× | 19.7 | 0.11 | 62 | A | 459 | 35.70 | 62 | A | 1.79 | 0.16 | 62 | AB | 1.25 | 0.12 | 62 | AB |
| Insecticide-1× | 19.8 | 0.11 | 62 | A | 393 | 36.70 | 62 | A | 1.62 | 0.16 | 62 | AB | 1.08 | 0.12 | 62 | AB |
| Insecticide-10× | 19.8 | 0.12 | 62 | A | 384 | 37.80 | 62 | A | 1.92 | 0.17 | 62 | AB | 1.28 | 0.12 | 62 | AB |
| Fungicide-1× | 19.7 | 0.11 | 62 | A | 402 | 36.20 | 62 | A | 2.17 | 0.16 | 62 | AB | 1.37 | 0.12 | 62 | AB |
| Fungicide-10× | 20.0 | 0.12 | 62 | A | 291 | 40.60 | 62 | A | 1.78 | 0.18 | 62 | AB | 1.04 | 0.13 | 62 | AB |

| 3a | 16S+ | | | | ITS+ | | | | AOA/AOB×100 | | | |
|---|---|---|---|---|---|---|---|---|---|---|---|---|
| | Estimate | SE | Z value | Pr(>\|z\|) | Estimate | SE | Z value | Pr(>\|z\|) | Estimate | SE | Z value | Pr(>\|z\|) |
| Herbicide-1× | -0.20 | 0.15 | -1.32 | 0.19 | 0.19 | 0.19 | 1.00 | 0.32 | -19.83 | 53.14 | -0.37 | 0.71 |
| Herbicide-10× | -0.14 | 0.13 | -1.05 | 0.29 | 0.07 | 0.17 | 0.44 | 0.66 | 96.26 | 46.48 | 2.07 | **0.04** |
| Mixture-1× | 0.11 | 0.14 | 0.77 | 0.44 | 0.40 | 0.18 | 2.19 | **0.03** | -38.66 | 50.62 | -0.76 | 0.45 |
| Mixture-10× | -0.07 | 0.15 | -0.44 | 0.66 | 0.27 | 0.19 | 1.44 | 0.15 | -56.62 | 51.77 | -1.09 | 0.27 |
| OM | 0.06 | 0.03 | 2.32 | **0.02** | 0.06 | 0.03 | 1.83 | 0.07 | 2.77 | 8.78 | 0.32 | 0.75 |
| pH | -1.41 | 0.66 | -2.13 | **0.03** | -2.29 | 0.83 | -2.76 | **0.01** | 289.21 | 232.48 | 1.24 | 0.21 |
| RATIOCN | -0.24 | 0.21 | -1.14 | 0.26 | -0.49 | 0.26 | -1.86 | 0.06 | 138.14 | 72.60 | 1.90 | 0.06 |
| CEC | 0.02 | 0.02 | 0.84 | 0.40 | 0.04 | 0.03 | 1.37 | 0.17 | -18.32 | 7.80 | -2.35 | **0.02** |

| 3b | AOB/16S×100 | | | | COMAA/16S×100 | | | | COMAB/16S×100 | | | |
|---|---|---|---|---|---|---|---|---|---|---|---|---|
| | Estimate | SE | Z value | Pr(>\|z\|) | Estimate | SE | Z value | Pr(>\|z\|) | Estimate | SE | Z value | Pr(>\|z\|) |
| Herbicide-1× | -0.04 | 0.13 | -0.31 | 0.75 | 0.97 | 0.21 | 4.72 | **<0.00** | 0.57 | 0.16 | 3.54 | **<0.01** |
| Herbicide-10× | -0.07 | 0.11 | -0.61 | 0.54 | 0.29 | 0.19 | 1.55 | 0.12 | 0.32 | 0.14 | 2.22 | **0.03** |
| Mixture-1× | 0.017 | 0.12 | 0.14 | 0.89 | 0.21 | 0.20 | 1.04 | 0.30 | 0.24 | 0.16 | 1.55 | 0.12 |
| Mixture-10× | 0.23 | 0.13 | 1.81 | 0.07 | 0.53 | 0.21 | 2.55 | **0.01** | 0.27 | 0.16 | 1.69 | 0.09 |
| OM | -0.03 | 0.02 | -1.42 | 0.16 | -0.04 | 0.04 | -1.05 | 0.29 | -0.03 | 0.03 | -1.10 | 0.27 |
| pH | -0.20 | 0.57 | -0.36 | 0.72 | -0.98 | 0.92 | -1.07 | 0.29 | -0.40 | 0.71 | -0.56 | 0.58 |
| RATIOCN | -0.30 | 0.18 | -1.68 | 0.09 | -0.49 | 0.29 | -1.71 | 0.09 | -0.14 | 0.22 | -0.61 | 0.54 |
| CEC | 0.06 | 0.02 | 3.19 | **<0.01** | -0.03 | 0.03 | -0.91 | 0.36 | -0.02 | 0.02 | -0.68 | 0.50 |

| Pairwise comparison-3 | ITS | | | | AOA/AOB×100 | | | | COMAA/16S×100 | | | | COMAB/16S×100 | | | |
|---|---|---|---|---|---|---|---|---|---|---|---|---|---|---|---|---|
| | Emmean | SE | df | group | Emmean | SE | df | group | Emmean | SE | df | group | Emmean | SE | df | group |
| Control | 19.1 | 0.12 | 33 | A | 377 | 33.70 | 32 | A | 1.38 | 0.14 | 33 | A | 0.88 | 0.10 | 33 | B |
| Herbicide-1× | 19.3 | 0.13 | 33 | A | 357 | 38.10 | 32 | A | 2.36 | 0.14 | 33 | B | 1.45 | 0.11 | 33 | A |
| Herbicide-10× | 19.2 | 0.13 | 33 | A | 474 | 34.80 | 32 | A | 1.67 | 0.14 | 33 | A | 1.20 | 0.11 | 33 | AB |
| Mixture-1× | 19.5 | 0.14 | 33 | A | 339 | 37.70 | 32 | A | 1.59 | 0.15 | 33 | A | 1.12 | 0.12 | 33 | AB |
| Mixture-10× | 19.4 | 0.14 | 33 | A | 321 | 37.60 | 32 | A | 1.91 | 0.15 | 33 | AB | 1.15 | 0.12 | 33 | AB |

| 4a | 16S+ | | | | ITS+ | | | | AOA/AOB×100 | | | |
|---|---|---|---|---|---|---|---|---|---|---|---|---|
| | Estimate | SE | Z value | Pr(>\|z\|) | Estimate | SE | Z value | Pr(>\|z\|) | Estimate | SE | Z value | Pr(>\|z\|) |
| Insecticide-1× | -0.12 | 0.16 | -0.78 | 0.44 | 0.12 | 0.15 | 0.83 | 0.40 | 12.50 | 59.60 | 0.21 | 0.83 |
| Insecticide-10× | -0.17 | 0.16 | -1.10 | 0.27 | 0.05 | 0.15 | 0.34 | 0.73 | 32.76 | 58.21 | 0.56 | 0.57 |
| Mixture-1× | 0.09 | 0.15 | 0.60 | 0.55 | 0.37 | 0.14 | 2.64 | **0.01** | 44.35 | 56.19 | 0.79 | 0.43 |
| Mixture-10× | 0.12 | 0.16 | 0.75 | 0.45 | 0.64 | 0.15 | 4.11 | **<0.01** | -78.59 | 61.62 | -1.28 | 0.20 |
| OM | 0.01 | 0.04 | 0.25 | 0.80 | -0.01 | 0.03 | -0.31 | 0.75 | 14.18 | 13.75 | 1.03 | 0.30 |
| pH | -0.42 | 0.62 | -0.68 | 0.50 | -0.39 | 0.59 | -0.67 | 0.51 | -69.88 | 234.45 | -0.30 | 0.77 |
| RATIOCN | -0.21 | 0.20 | -1.02 | 0.31 | -0.29 | 0.18 | -1.55 | 0.12 | 111.96 | 73.62 | 1.52 | 0.13 |
| CEC | 0.01 | 0.02 | 0.33 | 0.74 | 0.02 | 0.02 | 0.88 | 0.38 | 23.93 | 8.09 | -2.96 | **<0.01** |

| 4b | AOB/16S×100 | | | | COMAA/16S×100 | | | | COMAB/16S×100 | | | |
|---|---|---|---|---|---|---|---|---|---|---|---|---|
| | Estimate | SE | Z value | Pr(>\|z\|) | Estimate | SE | Z value | Pr(>\|z\|) | Estimate | SE | Z value | Pr(>\|z\|) |
| Insecticide-1× | -0.19 | 0.14 | -1.32 | 0.19 | 0.03 | 0.25 | 0.12 | 0.90 | 0.02 | 0.16 | 0.14 | 0.89 |
| Insecticide-10× | -0.29 | 0.14 | -2.13 | **0.03** | 0.57 | 0.25 | 2.30 | **0.02** | 0.30 | 0.16 | 1.90 | 0.06 |
| Mixture-1× | -0.20 | 0.13 | -1.52 | 0.13 | 0.57 | 0.24 | 2.41 | **0.02** | 0.23 | 0.15 | 1.48 | 0.14 |
| Mixture-10× | 0.14 | 0.15 | 0.96 | 0.34 | 0.32 | 0.26 | 1.23 | 0.22 | -0.09 | 0.17 | -0.51 | 0.61 |
| OM | -0.05 | 0.03 | -1.62 | 0.11 | 0.01 | 0.06 | 0.11 | 0.91 | 0.01 | 0.04 | 0.28 | 0.78 |
| pH | 0.72 | 0.56 | 1.30 | 0.19 | -1.29 | 0.99 | -1.29 | 0.20 | -0.09 | 0.64 | -0.15 | 0.88 |
| RATIOCN | -0.11 | 0.17 | -0.65 | 0.52 | -0.32 | 0.31 | -1.04 | 0.30 | 0.01 | 0.20 | 0.00 | 1.00 |
| CEC | 0.06 | 0.02 | 2.90 | **<0.01** | 0.05 | 0.03 | 1.48 | 0.14 | 0.01 | 0.02 | 0.13 | 0.90 |

| Pairwise comparison-4 | ITS | | | | AOB/16S×100 | | | | COMAA/16S×100 | | | |
|---|---|---|---|---|---|---|---|---|---|---|---|---|
| | Emmean | SE | df | group | Emmean | SE | df | group | Emmean | SE | df | group |
| Control | 19.1 | 0.10 | 34 | B | 0.94 | 0.09 | 34 | A | 1.49 | 0.17 | 34 | A |
| Insecticide-1× | 19.2 | 0.11 | 34 | B | 0.75 | 0.10 | 34 | A | 1.52 | 0.19 | 34 | A |
| Insecticide-10× | 19.1 | 0.11 | 34 | B | 0.64 | 0.11 | 34 | A | 2.05 | 0.19 | 34 | A |
| Mixture-1× | 19.5 | 0.10 | 34 | AB | 0.74 | 0.10 | 34 | A | 2.06 | 0.18 | 34 | A |
| Mixture-10× | 19.7 | 0.12 | 34 | A | 1.08 | 0.11 | 34 | A | 1.81 | 0.20 | 34 | A |

| 5a | 16S+ | | | | ITS+ | | | | AOA/AOB×100 | | | |
|---|---|---|---|---|---|---|---|---|---|---|---|---|
| | Estimate | SE | Z value | Pr(>\|z\|) | Estimate | SE | Z value | Pr(>\|z\|) | Estimate | SE | Z value | Pr(>\|z\|) |
| Fungicide-1× | 0.31 | 0.15 | -2.05 | **0.04** | -0.10 | 0.17 | -0.59 | 0.56 | 6.29 | 58.66 | 0.11 | 0.91 |
| Fungicide-10× | 0.09 | 0.18 | 0.49 | 0.62 | 0.35 | 0.20 | 1.75 | 0.08 | -112.85 | 68.62 | -1.65 | 0.10 |
| Mixture-1× | -0.57 | 0.15 | -3.81 | **<0.01** | -0.17 | 0.17 | -1.00 | 0.32 | 153.14 | 57.13 | 2.68 | **0.01** |
| Mixture-10× | -0.20 | 0.16 | -1.26 | 0.21 | 0.03 | 0.18 | 0.20 | 0.84 | -87.33 | 61.01 | -1.43 | 0.15 |
| OM | 0.03 | 0.04 | 0.78 | 0.43 | 0.02 | 0.04 | 0.63 | 0.53 | 3.20 | 13.89 | 0.23 | 0.82 |
| pH | 0.41 | 0.42 | 0.98 | 0.33 | 0.15 | 0.47 | 0.33 | 0.74 | -279.75 | 162.22 | -1.73 | 0.08 |
| RATIOCN | 0.03 | 0.15 | 0.23 | 0.82 | 0.17 | 0.17 | 0.98 | 0.33 | 13.53 | 58.60 | 0.23 | 0.82 |
| CEC | 0.03 | 0.02 | 1.53 | 0.13 | 0.05 | 0.02 | 2.03 | **0.04** | -22.27 | 8.04 | -2.77 | **0.01** |

| 5b | AOB/16S×100 | | | | COMAA/16S×100 | | | | COMAB/16S×100 | | | |
|---|---|---|---|---|---|---|---|---|---|---|---|---|
| | Estimate | SE | Z value | Pr(>\|z\|) | Estimate | SE | Z value | Pr(>\|z\|) | Estimate | SE | Z value | Pr(>\|z\|) |
| Fungicide-1× | 0.13 | 0.16 | 0.81 | 0.42 | 0.43 | 0.27 | 1.59 | 0.11 | 0.32 | 0.16 | 1.98 | **0.05** |
| Fungicide-10× | 0.42 | 0.19 | 2.28 | **0.02** | -0.05 | 0.32 | -0.16 | 0.87 | -0.04 | 0.19 | -0.21 | 0.84 |
| Mixture-1× | -0.16 | 0.16 | -1.04 | 0.30 | 0.56 | 0.26 | 2.11 | **0.03** | 0.31 | 0.16 | 1.96 | **0.05** |
| Mixture-10× | 0.35 | 0.17 | 2.09 | **0.04** | 0.29 | 0.28 | 1.02 | 0.31 | 0.20 | 0.17 | 1.18 | 0.24 |
| OM | -0.02 | 0.04 | -0.45 | 0.65 | -0.04 | 0.06 | -0.62 | 0.54 | -0.02 | 0.04 | -0.45 | 0.65 |
| pH | 1.14 | 0.44 | 2.58 | **0.01** | -0.53 | 0.75 | -0.71 | 0.48 | -0.05 | 0.45 | -0.11 | 0.92 |
| RATIOCN | -0.04 | 0.16 | -0.28 | 0.78 | 0.13 | 0.27 | 0.50 | 0.62 | -0.01 | 0.16 | -0.08 | 0.94 |
| CEC | 0.02 | 0.02 | 1.02 | 0.31 | 0.04 | 0.04 | 0.97 | 0.33 | 0.01 | 0.02 | 0.20 | 0.85 |

| Pairwise comparison-5a | 16S | | | | AOA/AOB×100 | | | | AOB/16S×100 | | | |
|---|---|---|---|---|---|---|---|---|---|---|---|---|
| | Emmean | SE | df | group | Emmean | SE | df | group | Emmean | SE | df | group |
| Control | 20.0 | 0.11 | 34 | A | 382 | 41.60 | 34 | AB | 0.74 | 0.11 | 34 | AB |
| Fungicide-1× | 19.7 | 0.11 | 34 | AB | 388 | 40.70 | 34 | AB | 0.86 | 0.11 | 34 | AB |
| Fungicide-10× | 20.1 | 0.13 | 34 | A | 269 | 48.20 | 34 | B | 1.16 | 0.13 | 34 | A |
| Mixture-1× | 19.4 | 0.12 | 34 | B | 535 | 45.30 | 34 | A | 0.57 | 0.12 | 34 | B |
| Mixture-10× | 19.8 | 0.11 | 34 | AB | 294 | 42.20 | 34 | B | 1.08 | 0.12 | 34 | AB |

| Pairwise comparison-5b | COMAA/16S×100 | | | | COMAB/16S×100 | | | |
|---|---|---|---|---|---|---|---|---|
| | Emmean | SE | df | group | Emmean | SE | df | group |
| Control | 1.70 | 0.20 | 34 | A | 1.03 | 0.12 | 34 | A |
| Fungicide-1× | 2.13 | 0.20 | 34 | A | 1.35 | 0.12 | 34 | A |
| Fungicide-10× | 1.65 | 0.23 | 34 | A | 0.99 | 0.14 | 34 | A |
| Mixture-1× | 2.26 | 0.22 | 34 | A | 1.34 | 0.13 | 34 | A |
| Mixture-10× | 1.99 | 0.20 | 34 | A | 1.23 | 0.12 | 34 | A |

Table S4.

| 1 | Nematode abundance | | | |
|---|---|---|---|---|
| | Estimate | SE | Z value | Pr(>\|z\|) |
| Mixture-1× | -222.46 | 71.43 | -3.11 | **<0.01** |
| Mixture-10× | 46.64 | 79.24 | 0.59 | 0.56 |
| OM | -26.31 | 19.68 | -1.34 | 0.18 |
| pH | 783.08 | 427.73 | 1.83 | 0.07 |
| RATIOCN | 149.74 | 127.26 | 1.18 | 0.24 |
| CEC | 41.62 | 13.76 | 3.03 | **<0.01** |

| Pairwise comparison-1 | Nematode abundance | | | |
|---|---|---|---|---|
| | Emmean | SE | df | group |
| Control | 1132 | 48.30 | 54 | A |
| Mixture-1× | 910 | 53.70 | 54 | B |
| Mixture-10× | 1179 | 57.20 | 54 | A |

| 2 | Nematode abundance | | | |
|---|---|---|---|---|
| | Estimate | SE | Z value | Pr(>\|z\|) |
| Herbicide-1× | 189.21 | 113.84 | 1.66 | 0.10 |
| Herbicide-10× | -56.80 | 109.44 | -0.52 | 0.60 |
| Insecticide-1× | -125.90 | 114.37 | -1.10 | 0.27 |
| Insecticide-10× | -138.46 | 108.93 | -1.27 | 0.20 |
| Fungicide-1× | -150.04 | 114.12 | -1.32 | 0.19 |
| Fungicide-10× | 235.08 | 125.20 | 1.88 | 0.06 |
| OM | 14.54 | 17.47 | 0.83 | 0.41 |
| pH | 507.05 | 315.37 | 1.61 | 0.11 |
| RATIOCN | -62.82 | 131.05 | -0.48 | 0.63 |
| CEC | 47.16 | 12.99 | 3.63 | **<0.01** |

| 3 | Nematode abundance | | | |
|---|---|---|---|---|
| | Estimate | SE | Z value | Pr(>\|z\|) |
| Herbicide-1× | 187.71 | 128.62 | 1.46 | 0.14 |
| Herbicide-10× | -8.81 | 115.98 | -0.08 | 0.94 |
| Mixture-1× | -121.03 | 124.23 | -0.97 | 0.33 |
| Mixture-10× | 53.55 | 128.99 | 0.42 | 0.68 |
| OM | 28.79 | 21.85 | 1.32 | 0.19 |
| pH | 459.87 | 570.54 | 0.81 | 0.42 |
| RATIOCN | -50.78 | 180.78 | -0.28 | 0.78 |
| CEC | 51.59 | 19.21 | 2.69 | **0.01** |

| 4 | Nematode abundance | | | |
|---|---|---|---|---|
| | Estimate | SE | Z value | Pr(>\|z\|) |
| Insecticide-1× | -99.67 | 94.72 | -1.05 | 0.29 |
| Insecticide-10× | -218.19 | 92.50 | -2.36 | **0.02** |
| Mixture-1× | -204.08 | 89.29 | -2.29 | **0.02** |
| Mixture-10× | 235.20 | 97.93 | 2.40 | **0.02** |
| OM | -46.28 | 21.85 | -2.12 | **0.03** |
| pH | 966.29 | 372.59 | 2.59 | **0.01** |
| RATIOCN | 95.99 | 116.99 | 0.82 | 0.41 |
| CEC | 10.30 | 12.86 | 0.80 | 0.42 |

| Pairwise comparison-4 | Nematode abundance | | | |
|---|---|---|---|---|
| | Emmean | SE | df | group |
| Control | 1073 | 75.70 | 34 | AB |
| Insecticide-1× | 974 | 81.80 | 34 | B |
| Insecticide-10× | 855 | 82.70 | 34 | B |
| Mixture-1× | 869 | 78.50 | 34 | B |
| Mixture-10× | 1309 | 85.30 | 34 | A |

| 5 | Nematode abundance | | | |
|---|---|---|---|---|
| | Estimate | SE | Z value | Pr(>\|z\|) |
| Fungicide-1× | -176.62 | 123.43 | -1.43 | 0.15 |
| Fungicide-10× | 283.79 | 144.38 | 1.97 | **0.05** |
| Mixture-1× | -247.05 | 120.20 | -2.06 | **0.04** |
| Mixture-10× | 45.40 | 128.38 | 0.35 | 0.72 |
| OM | -33.56 | 29.23 | -1.15 | 0.25 |
| pH | 610.19 | 341.31 | 1.79 | 0.07 |
| RATIOCN | 212.23 | 123.30 | 1.72 | 0.09 |
| CEC | 48.59 | 16.91 | 2.87 | **<0.01** |

| Pairwise comparison-5 | Nematode abundance | | | |
|---|---|---|---|---|
| | Emmean | SE | df | group |
| Control | 1119 | 87.50 | 34 | AB |
| Fungicide-1× | 943 | 85.70 | 34 | B |
| Fungicide-10× | 1403 | 101.50 | 34 | A |
| Mixture-1× | 872 | 95.30 | 34 | B |
| Mixture-10× | 1165 | 88.70 | 34 | AB |

Table S5.

| Analysis | | Factor | Df | Mean sq | F-value | Pr(>F) |
|---|---|---|---|---|---|---|
| Herbicide | Short-term | Treatment-Dose | 2 | 0.004126 | 0.268 | 0.773 |
| | Long-term | | | 0.00606 | 0.16 | 0.856 |
| Insecticide | Short-term | Treatment-Dose | 2 | 0.01880 | 0.66 | 0.551 |
| | Long-term | | | 0.01649 | 0.495 | 0.633 |
| Fungicide | Short-term | Treatment-Dose | 2 | 0.03003 | 0.487 | 0.637 |
| | Long-term | | | 0.01886 | 0.338 | 0.726 |
| Mixture | Short-term | Treatment-Dose | 2 | 0.06136 | 4.882 | 0.0551 |
| | Long-term | | | 0.03876 | 1.453 | 0.306 |
| | Short×long term | Treatment-Dose | 2 | 0.08828 | 4.499 | **0.0348** |
| | | Ratio | 1 | 0.00084 | 0.043 | 0.8397 |
| | | Treatment-Dose×Ratio | 2 | 0.02367 | 0.603 | 0.5629 |

Table S6.

| | | Herbicide | | | Insecticide | | | Fungicide | | | Mixture | | |
|---|---|---|---|---|---|---|---|---|---|---|---|---|---|
| Dose | | 1x | 10x | Control | 1x | 10x | Control | 1x | 10x | Control | 1x | 10x | Control |
| Ratio | Short term | -0.01±0.14 | 0.05±0.12 | 0.04±0.10 | 0.11±0.26 | 0.04±0.11 | -0.05±0.09 | 0.29±0.41 | 0.09±0.09 | 0.17±0.03 | -0.11±0.05 | -0.04±0.18 | 0.16±0.05 |
| | Long term | 0.07±0.24 | 0.04±0.23 | 0.13±0.05 | 0.23±0.30 | 0.08±0.08 | 0.14±0.04 | 0.17±0.37 | 0.01±0.03 | 0.11±0.16 | -0.11±0.20 | 0.07±0.07 | 0.10±0.18 |

Table S1. Results of the GLMM models and pairwise comparisons on the bacteria alpha diversity. Significant difference is indicated in bold

Table S2. Results of the GLMM models and pairwise comparisons on the fungi alpha diversity. Significant difference is indicated in bold

Table S3. Results of the GLMM models and pairwise comparisons on the microbial abundance parameters. Significant difference is indicated in bold. + indicates log transformation of the data

Table S4. Results of the GLMM models and pairwise comparisons on the nematode abundance. Significant difference is indicated in bold

Table S5. Analysis of variance (ANOVA) result of the short term or long-term (ratio) effect of the treatments (herbicide, insecticide, fungicide and mixture) on the nematode abundance, expressed as log10-transformed ratio between the post-application nematode abundances at day 7 or day 28, by the initial abundance (day -1). Significant differences (p=0.05) are indicated in bold

Table S6. Summary (mean and standard deviation) of the short term or long-term (ratio) effect of the treatments (herbicide, insecticide, fungicide and mixture) on the nematode abundance, expressed as log10-transformed ratio between the post-application nematode abundances at day 7 or day 28, by the initial abundance (day -1)
